# Cell Type Differentiation Using Network Clustering Algorithms

**DOI:** 10.1101/2024.12.04.626793

**Authors:** Fatemeh Sadat Fatemi Nasrollahi, Filipi Nascimento Silva, Shiwei Liu, Soumilee Chaudhuri, Meichen Yu, Juexin Wang, Kwangsik Nho, Andrew J. Saykin, David A. Bennett, Olaf Sporns, Santo Fortunato

**Affiliations:** Observatory of Social Media, Luddy School of Informatics, Computing, and Engineering, Indiana University, Indiana, USA; Center for Neuroimaging and the Indiana Alzheimer’s Disease Research Center, Indiana University, Indiana, USA; Department of Biomedical Engineering and Informatics, Luddy School of Informatics, Computing, and Engineering, Indiana University, Indiana, USA; Department of Psychology, Indiana University, Indiana, USA; Rush Alzheimer’s Disease Center (Drs. Bennett, Schneider, and Wilson) and Rush Institute for Healthy Aging (Drs. Bienias and Evans), Rush University Medical Center, Illinois, USA

**Keywords:** single-cell RNA-seq, cell separation, network clustering

## Abstract

Single cell RNA-seq (scRNA-seq) technologies provide unprecedented resolution representing transcriptomics at the level of single cell. One of the biggest challenges in scRNA-seq data analysis is the cell type annotation, which is usually inferred by cell separation approaches. In-silico algorithms that accurately identify individual cell types in ongoing single-cell sequencing studies are crucial for unlocking cellular heterogeneity and understanding the biological basis of diseases. In this study, we focus on robustly identifying cell types in single-cell RNA sequencing data; we conduct a comparative analysis using methods established in biology, like Seurat, Leiden, and WGCNA, as well as Infomap, statistical inference via Stochastic Block Models (SBM), and single-cell Graph Neural Networks (scGNN). We also analyze preprocessing pipelines to identify and optimize key components in the process. Leveraging two independent datasets, PBMC and ROSMAP, we employ clustering algorithms on cell-cell networks derived from gene expression data. Our findings reveal that while clusters detected by WGCNA exhibit limited correspondence with cell types, those identified by multiresolution Infomap and Leiden, and SBM show a closer alignment, with Infomap standing out as a particularly effective approach. Infomap notably offers valuable insights for the precise characterization of cellular landscapes related to neurodegenration and immunology in scRNA-seq.

## 1. Introduction

In recent years, significant advances in single-cell RNA sequencing (scRNA-Seq) technologies have revolutionized our ability to dissect cellular heterogeneity and uncover deep biological insights [1, 2, 3, 4, 5]. Unlike traditional bulk gene expression analysis, which provides average information from a population of cells, scRNA-Seq offers a granular view by quantifying mRNA expression in individual cells. This approach not only facilitates the identification of distinct cell types, it also enables the characterization of cell-specific gene expression profiles and functions. The workflow of single-cell sequencing typically involves isolating individual cells using sophisticated techniques such as Fluorescence-Activated Cell Sorting (FACS) and Droplet-Based Technologies like the 10× Genomics Chromium system. Droplet-based methods have gained popularity for their ability to simultaneously analyze thousands of cells, leveraging Unique Molecular Identifiers (UMIs) to minimize Polymerase Chain Reaction (PCR) bias and accurately annotate transcripts [2, 4, 5].

Single-cell studies encompass a wide array of biological and medical inquiries, ranging from delineating cell lineage and identifying novel cell type marker genes to predicting cell fate and exploring gene expression dynamics in disease contexts [6, 7, 8, 9]. To achieve these objectives, researchers often employ diverse clustering methods tailored to uncover meaningful patterns within scRNA-Seq datasets [10, 11, 12, 13]. These methods streamline the identification of both well-characterized and novel cell types, providing crucial insights into cellular function and behavior under physiological and pathological conditions.

Clustering results vary significantly across different methods, aligning with previous findings [14, 15]. Such variation is expected, as clustering methods differ in key aspects, like the concept of the cluster itself. Each clustering algorithm has its unique advantages and drawbacks. Therefore, employing multiple clustering methods can provide a comprehensive understanding of cell clusters from different perspectives and a platform to compare the methods.

One significant challenge in scRNA-seq analysis is the need for users to carefully select clustering parameters that provide the most accurate approximation of the true biological clusters. Many clustering methods typically involve multiple parameters, and determining the optimal settings is not a trivial task. The complexity of this parameter selection process can significantly impact the performance and reliability of the outcomes. On the other hand, it is desirable that results do not depend too much on specific parameter choices or remain stable over sufficiently large parameter ranges. Additionally, addressing these challenges necessitates the development of tools that facilitate data analysis and parameter exploration. In particular, interactive visualization interfaces can play a crucial role in enhancing the accessibility and interpretability of clustering analyses in bioinformatics [16]. Such interfaces enable researchers to intuitively explore parameter spaces, assess the stability of clustering results, and make more informed decisions regarding parameter selection.

Certain packages, such as Seurat [17, 18], are widely used for clustering within the scRNA-seq data analysis. Despite their widespread use in various applications, there remains substantial potential for enhancing cell clustering analysis. Previous studies have systematically compared outcomes from multiple methods, including Seurat and Weighted Gene Co-expression Network Analysis (WGCNA) [19]. Many methods predominantly utilize hierarchical clustering and k-means as their clustering strategies [20, 21, 22]; however, these techniques have notable limitations. Hierarchical clustering can be computationally demanding, especially with large datasets, and often struggles to identify optimal cluster structures when the data does not naturally follow a hierarchical pattern. Additionally, it typically lacks a clear criterion for selecting meaningful clusters from the many partitions it produces. K-means clustering, on the other hand, requires the number of clusters to be specified in advance — an unknown factor in most cases — and assumes clusters are convex in shape, which may misrepresent the true structure of the data.

Therefore, there is a critical need to identify and develop accurate algorithms that surpass such limitations. Novel algorithms, proposed and refined in other fields, have shown great promise but are often underutilized in scRNA-seq analysis. By systematically evaluating and adapting these cutting-edge algorithms, we can significantly advance the precision and effectiveness of clustering analyses in biological research.

Preprocessing is another critical component of the analysis, as it transforms raw data into networks, which clustering algorithms can then interpret. This is particularly important in scRNA-seq analysis due to complexity, noise, and high dimensionality of biological data. Effective preprocessing usually consists of multiple steps: normalization to correct for technical variations, noise reduction to minimize random fluctuations, feature selection to highlight the most informative genes, and dimension reduction to simplify the data while preserving key information. These steps are essential for improving data quality and uncovering biological signals. However, the preprocessing steps in different packages are often complex and challenging for users to implement correctly. Therefore, it is important to thoroughly examine these preprocessing steps in the context of scRNA-seq data and comprehensively report the effects of each.

In this study, we approach the identification of cell types as a network clustering problem [23, 24, 25]. We construct networks from gene expression data derived from two distinct scRNA-seq datasets and apply several state-of-the-art clustering algorithms, including some that are rarely used in biology. Our contribution is twofold: first, to identify the best-performing algorithms for cluster identification, using a reliable ground truth; second, to propose a clear and user-friendly preprocessing pipeline that can be easily tuned for each dataset to achieve optimal results. This work advances the precision and reliability of clustering analyses in biological research. We believe that this study not only contributes to increasing the accuracy of cell type identification, it also provides researchers with a more accessible and effective toolkit for handling complex biological datasets.

## 2. Materials and Methods

### 2.1. Clustering algorithms for scRNA-seq analysis

In this section, we provide a brief review of the network clustering algorithms employed in this study. These include Leiden [26], Infomap [27, 28], WGCNA [29], statistical inference through the Stochastic Block Model (SBM) [30], Seurat [17, 18], and single-cell Graph Neural Network (scGNN) [31].

**Leiden** optimizes modularity, a quality function that rewards partitions with higher density of edges within clusters than expected in a randomized network with the same degree sequence of the input one [26, 32]. The resolution parameter in the Leiden algorithm allows one to adjust the level of granularity of the analysis, going from small to large clusters, as required by the biological context of the dataset. Moreover, Leiden can operate on both weighted and unweighted networks. Particularly with the incorporation of edge weights, the Leiden algorithm can detect cell clusters with nuanced transcriptional variances.

**Infomap** is designed to identify clusters in networks by optimizing the compression of the information used to describe a diffusion process running on the network [27]. Infomap includes a parameter known as the Markov time [33], which governs the pace of the exploration of the network’s structure via the diffusion process and influences the granularity of clustering. Similarly to changing the resolution parameter in Leiden, adjusting the Markov time enables analysts to fine-tune the clustering resolution according to the desired level of detail in the analysis. Infomap is capable of analyzing both weighted and unweighted networks.

**WGCNA (Weighted Gene Co-expression Network Analysis)** analyzes correlation patterns in gene expression data across multiple samples. Typically, it constructs a weighted gene co-expression network to identify modules of highly correlated genes associated with specific biological functions or traits using hierarchical clustering [34, 29]. WGCNA is widely used in the analysis of regulatory networks and gene co-expression relationships in complex biological systems. In this work, we adapted WGCNA to cluster cells instead of genes. By analyzing gene expression profiles across cells, WGCNA constructs a cell co-expression network and identifies clusters of cells with similar expression patterns. In this study, due to the considerable size of the gene space, the conventional approach of constructing a cell co-expression network directly encountered significant challenges. To address this, Principal Component Analysis (PCA) was employed to effectively reduce the dimensionality of the dataset. Specifically, we use 50 principal components in the study, which is the default setting in Seurat, to maintain consistency.

**Statistical inference via Stochastic Block Model (SBM)**. The Stochastic Block Model is a probabilistic generative model of networks with community structure [35]. Its distinctive feature is that the probability that two nodes are connected solely depends on their community memberships. By fitting an SBM to a network, one then recovers the partition upon which the model generates a network most similar to the input one. Nested SBM derives a hierarchical partition, with clusters including (included in) smaller (larger) clusters [36, 30]. Such a hierarchical approach facilitates the identification of cluster structures at different resolutions, as the different levels of the hierarchy correspond to different cluster sizes. In contrast to multiresolution methods like Leiden and Infomap, the nested SBM lacks a resolution parameter, offering only a limited number of possible divisions. Furthermore, a key drawback of the nested SBM is its significantly higher time complexity compared to both Leiden and Infomap.

**Seurat** provides built-in functions for clustering scRNA-seq data [17, 18]. These functions enable the partitioning of cells into clusters based on gene expression similarity or network connectivity. Seurat uses Louvain modularity Optimizer by [37] as the default modularity maximization method. Since the Leiden algorithm is just a modification of Louvain, the clustering outcomes from Seurat are expected to exhibit concordance with the partitions derived from Leiden. Throughout this work, the term “Seurat” specifically refers to the Seurat framework utilizing the Louvain modularity maximization algorithm.

**scGNN (Single-cell Graph Neural Network)** stacks graph neural networks in an iterative framework tailored for scRNA-seq analysis [31]. It leverages graph neural networks to capture complex relationships in cell similarity networks. scGNN learns low-dimensional embeddings for cells, facilitating clustering and imputation tasks while preserving biological relevance. scGNN considers unweighted interactions in the construction of these networks.

### 2.2. Data sets

We used a well-characterized 68k human Peripheral Blood Mononuclear Cells (PBMCs) scRNA-seq dataset via the 10× Genomics Chromium platform [5, 38, 4]. The PBMC dataset comprises single-cell transcriptomes from 68k freshly isolated PBMCs. PBMCs represent a heterogeneous population of white blood cells in peripheral blood, consisting of mostly lymphocytes (T cells, B cells, and natural killer (NK) cells), monocytes, and dendritic cells. Cell types were assigned by first down-sampling each purified PBMC population to 16,000 confidently mapped reads per cell. Then, the average gene expression profile was calculated for each population. The gene expression of each cell in the complex population was compared to these profiles using Spearman’s correlation. Each cell was assigned the ID of the purified population with which it had the highest correlation.

The PBMC dataset is notable for its balance between cell count and the features provided by droplet-based technology. The diverse array of profiled cells supports a detailed characterization of immune cell diversity, enabling the detection of less abundant cell populations that might be challenging to identify in smaller datasets. Similarly to the works of Sun et al. [39] and Huh et al. [19], we sampled the data as follows:

1. 20k simple case: This subset comprises all 20k cells from three distinctly discernible cell types: CD56+NK cells, CD19+B cells, and CD4+/CD25+ regulatory T cells.
2. 2k challenging case: This subset includes 2k cells from three closely related cell types with similar expression profiles: CD8+/CD45RA+ naive cytotoxic T cells, CD4+/CD25+ regulatory T cells, and CD4+/CD45RA+/CD25-naive T cells.

Each of these subsets, along with the full dataset of 68k PBMCs, was used as a separate input to conduct a comparative analysis of the clustering algorithms’ performance across different subsets.

Furthermore, we utilized the ROSMAP single-cell RNA sequencing dataset, derived from the dorsolateral prefrontal cortex (DLPFC) region of 12 male and 12 female participants, as described by Cain et al. [40]. ROSMAP stands for Religious Orders Study (ROS)/ Rush Memory and Aging Project (MAP); ROSMAP enroll older persons without known dementia all of whom agreed to annual clinical evaluation and brain donation at death [41]. Both studies were approved by an Institutional Review Board of Rush University Medical Center. All participants signed informed and repository consents and an Anatomic Gift Act. The comprehensive dataset used here comprises 172,659 cells, including astrocytes, excitatory neurons, inhibitory neurons, oligodendrocytes, microglia, endothelial cells, and pericytes, which were annotated using unsupervised clustering algorithms in the aforementioned study. The processed single-cell data and corresponding cell type annotations were sourced from Synapse (syn16780177).

This dataset consists of cells annotated by the Seurat pipeline that expressed marker genes from multiple cell types, such as endothelial cells and pericytes, indicating the presence of doublet cells that required further refinement. We employed Azimuth [42] as another reference and retained cells that exhibited consistent cell type annotations from both the Cain et al. study and the Azimuth prediction, thereby ensuring a high level of annotation accuracy and reliability in our subsequent analyses; the filtered dataset consists of about 21k cells. Figure 1 represents the expression levels of several well-known brain marker genes [43, 44, 10] across the types assigned to cells in this dataset. One observes that there is a strong correspondence between the marker gene expression levels and cell types, which underscores the reliability of the cell type annotations in this filtered dataset.

**Fig. 1.**
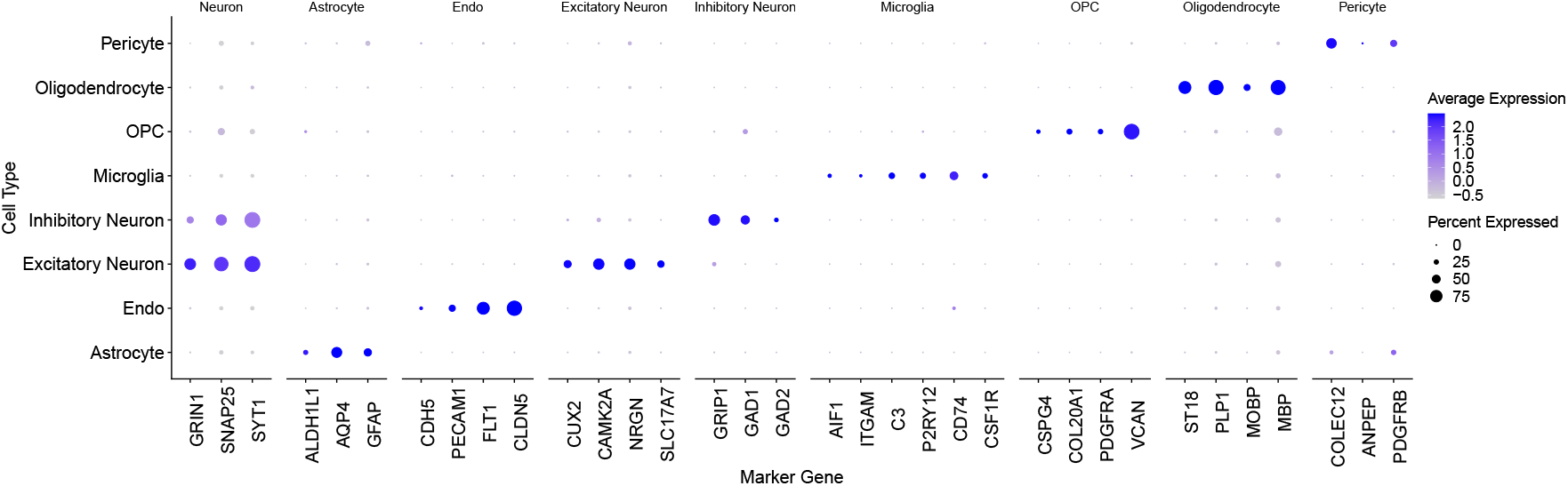
Expression levels of brain cell marker genes across the cell types in the ROSMAP dataset. The size of each dot represents the proportion of cells in a particular cluster or cell type that express a given gene. A larger dot indicates that a higher percentage of cells in that cluster express the gene. The color of each dot reflects the average expression level of the gene in the cells that express it; darker blue indicates higher expression levels.

### 2.3. Preprocessing techniques

#### 2.3.1. Preprocessing pipeline using Seurat R package

In the preprocessing pipeline for scRNA-seq data using the Seurat package in R [18], several essential steps are followed as seen in Figure 2. Initially, a normalization function `NormalizeData()` is applied to correct technical biases and normalize expression levels across cells, essential for ensuring comparability during downstream analysis. Identifying genes that exhibit high variability across cells is crucial for capturing informative features: this is achieved through the `FindVariableFeatures()` function. Scaling the data with `ScaleData()` standardizes expression values, preventing genes with larger expressions from dominating subsequent analyses. Dimensionality reduction is then performed using PCA with `RunPCA()`, extracting the major sources of variation in the dataset. Finally, FindNeighbors() calculates cell similarities, laying the groundwork for tasks such as clustering and visualization. K-nearest neighbors (KNN) and shared nearest neighbors (SNN) algorithms are employed; through KNN, each cell identifies its closest neighbors based on similarity metrics, constructing a weighted undirected network where cells are connected to their most similar counterparts. Additionally, SNN evaluates shared neighbors between cells, emphasizing connections that go beyond immediate proximity and capture higher-order relationships. By leveraging these network-based approaches, this technique transforms the single-cell expression data into a network representation that encapsulates complex cellular interactions and structural properties. This network is then used for the clustering analysis, which partition cells into biologically meaningful groups based on their network connectivity patterns. By integrating network-derived insights into the clustering process, this approach enhances the resolution and accuracy of cell type identification and characterization, enabling comprehensive exploration of cellular heterogeneity and dynamics.

**Fig. 2.**
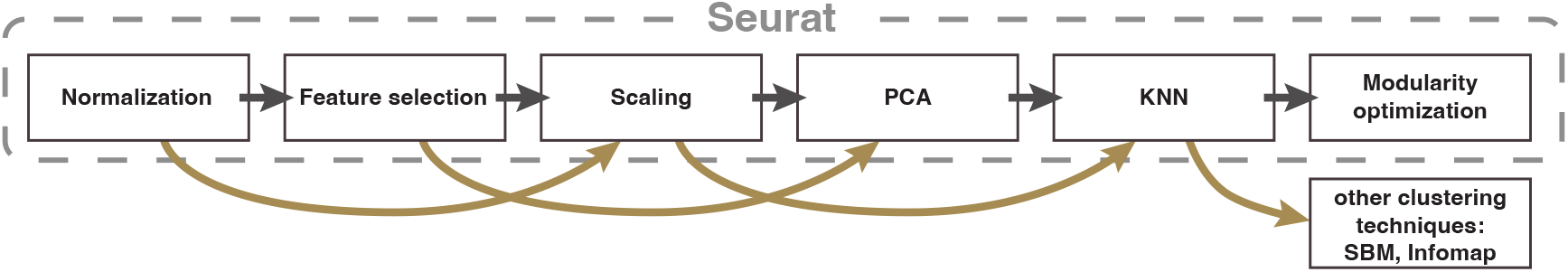
Schematic representation of the methodology. The Seurat pipeline includes all steps up to network clustering via modularity maximization. The alternative pipeline allows for flexibility by enabling exploration of the Seurat pipeline steps, with the option to skip one or more steps until the KNN network is generated (paths shown in yellow). Additionally, the alternative pipeline offers the choice of using clustering methods beyond modularity maximization, such as SBM and Infomap.

#### 2.3.2. Alternative preprocessing pipeline

The process of constructing biological networks from diverse datasets is a complex task as it encompasses a wide range of methodologies and critical decisions. These choices can significantly influence the network representation, and as a consequence, impacts cluster detection. In functional brain networks, one common approach involves the calculation of correlation coefficients (either Pearson or Spearman) across the time series of functional Magnetic Resonance Imaging(fMRI) or electroencephalogram (EEG) signals [45]. This method facilitates the identification of synchronous activity patterns among brain regions, where nodes represent distinct areas of the brain. Similarly, gene co-expression networks are derived by calculating the correlation between transcriptomes for each gene pair across a population of individuals. In these networks, genes are represented as nodes, typically numbering between 10,000 to 20,000.

A common preprocessing pipeline may involve several steps, such as transformation or normalization [46, 47], feature selection [42], scaling and dimension reduction [48]. The normalization ensures that the gene expression values are comparable across cells, and can be done by dividing the counts for each cell by the total count. In addition, the data can be log-transformed (using log(count + 1)), which reduces the effects of outliers and stabilizes the variance. Data can then be centered and scaled through standardization, which involves computing the z-scores of the values based on the mean and standard deviation of the null model distribution. In this case, the null model distribution is derived from an estimated mean-variance relationship, obtained by fitting a second-degree polynomial to the variance-vs-mean curve across all gene entries. The standardization procedure adjust each gene entry by subtracting its mean and dividing by the estimated standard deviation derived from the polynomial fit. Next, features are selected to reduce noise in the data and better encode genes that capture significant biological variability.

Since the number of genes in scRNA-seq can be large, and many gene profiles may correlate to each other, dimension reduction can be used to reduce the computational cost of network construction, while maintaining the relevant relationships between cells. A common approach is PCA [48], which reduces the data to linear combinations of maximum variance.

The network of cells can be constructed through several approaches. The most common is computing the similarity between all pairs of cells. However such a procedure does not scale well with the size of the networks. Alternatively, one can use KNN [46]; both approaches use Pearson’s correlation between the cell profiles across genes. It is generally unfeasible to calculate, store, and analyze all pairwise similarities, particularly for large datasets. For that reason, sparsification strategies are normally employed, like removing all similarities lower than a fixed threshold, or using localized methods, such as the disparity filter [49]. For very large datasets, neither of these options may be suitable given their computational costs and required memory to run. In that case, approximated approaches for KNN can be used, such as Nearest Neighbor Descent [50].

While all the aforementioned steps of the preprocessing pipeline have been used in most procedures for network generation, it is unclear how they affect the network structure and, consequently, the detected clusters. Here, we investigate how different choices for the pipeline demonstrated in Figure 2 may lead to different clustering performances.

## 3. Results

### 3.1. Results on PBMCs: Seurat-generated networks

In this section we present the results of applying the six clustering algorithms described in section Clustering algorithms for scRNA-seq analysis to the networks obtained from the PBMCs data (see section 2.2). The networks analyzed in this section are generated using the Seurat pipeline as described in section 2.3.1. Clustering analysis can be conducted with or without considering edge weights produced by Seurat, allowing for a comparison of clustering algorithms’ performance in both the weighted and the unweighted case. As described in section 2.2, there are two subsets of the PBMCs data considered in this analysis. The full data set is also used to generate a network of 68k nodes, representing cells, and edges, representing cell-cell similarities.

The similarity between the detected clusters and the cell types is estimated via the Adjusted Rand Index (ARI) [51], which reaches a maximum of 1 when the partitions are identical, and around 0 if they are randomly correlated.

First, we consider the 20k dataset, which comprises cells from highly distinct cell types. Figure 3 (left) illustrates the ARI for each method.

**Fig. 3.**
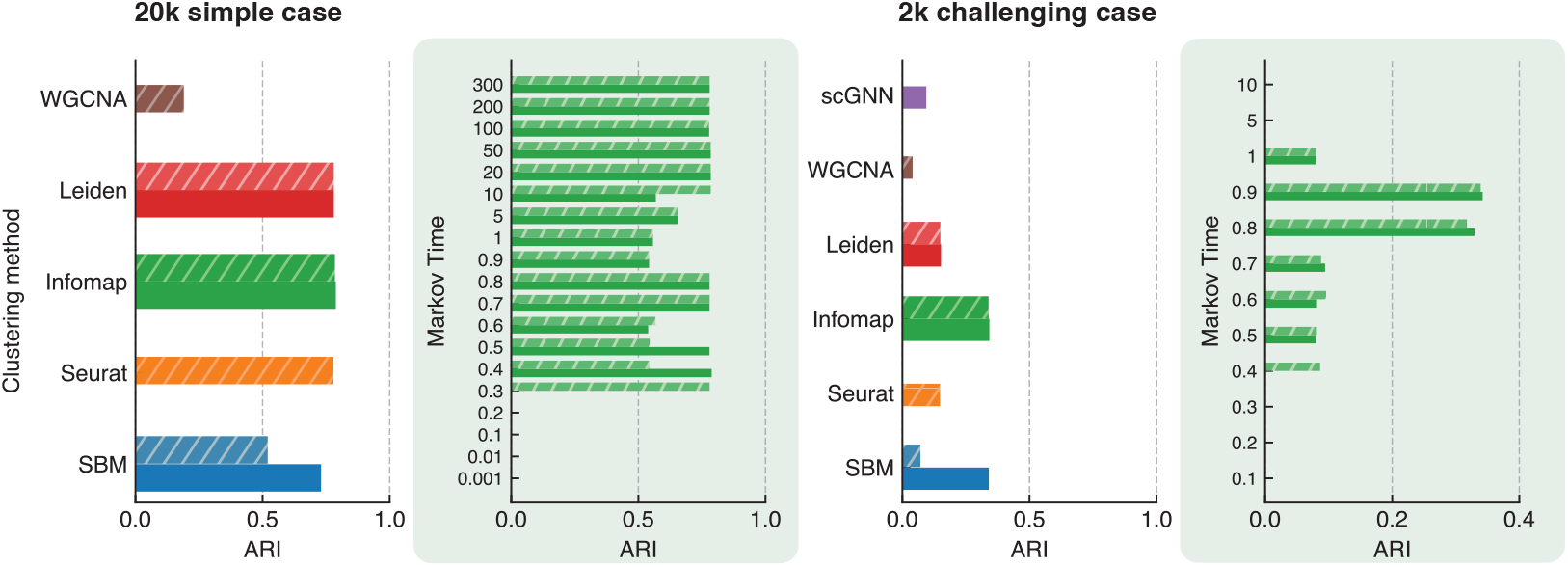
Adjusted Rand Index (ARI) between cell types and detected clusters for SBM, Seurat, Infomap, Leiden, WGCNA, and scGNN in the 20k simple (left panel) and 2k challenging PBMCs datasets (right panel). scGNN is not included in the left panel due to its high time complexity. Both the weighted (hatched) and unweighted (unhatched) versions of the same networks were considered for algorithms that can handle both. The zoomed-in panels in green illustrate the ARI for Infomap across different Markov times.

In this study, the Leiden and Infomap algorithms were executed 100 times each. For Leiden, the partition with the highest modularity was selected, while for Infomap, the partition with the lowest description length was chosen. Due to the higher computational demands of SBM, it was run 20 times, and the partition with the lowest description length was used for the ARI calculation. In contrast, Seurat, WGCNA, and scGNN were each run once, and their respective partitions were directly used to compute the ARI.

Given the clear distinction between the cell types in this dataset (see Figure S1, all algorithms are expected to perform relatively well. This expectation is confirmed in Figure 3 left panel, where the ARI for Seurat, Infomap, and Leiden is 0.78 and for SBM is 0.73. WGCNA performs poorly, with an ARI of 0.19 on this dataset, potentially due to its requirement for constructing a scale-free network [52] based on the data. In this case, a KNN-based network appears to be more effective than the scale-free approach. The zoomed-in figure in green details the ARI for Infomap and different Markov times, highlighting its stable and robust performance across a broad range of Markov times, which makes it the most reliable method in this context. For comparison with other methods across different resolution parameters, see Figure S2. Note that scGNN is computationally expensive for this dataset due to its large size and thus it is not included in this figure.

To rigorously evaluate and compare the performance of the algorithms on more challenging cell types — those with similar gene expression profiles — we extracted a representative sample of 2,000 cells. This sample was randomly selected while maintaining the same proportional distribution of cell types as in the original dataset. Figure 3 right panel shows the results of this analysis. This dataset presents significant challenges for all clustering methods (see Figure S3). The highest ARI achieved is 0.34, for SBM and Infomap. Seurat and Leiden perform less effectively, with an ARI of 0.15. Notably, WGCNA and scGNN exhibit poor performance on this dataset, as illustrated in the figure. The zoomed-in figure in green highlights the same challenge when applying Infomap to this dataset, showing that its performance stability is confined to a narrower range of Markov times. Nevertheless, when compared with other methods across various resolution parameters in Figure S4, Infomap has a superior performance overall.

Finally, we applied the algorithms to the complete dataset, encompassing all 68k PBMCs. Figure 4 reports the ARI for the methods that were able to complete the analysis in a reasonable time interval. The ARIs achieved are 0.35, 0.34, 0.30, and 0.27, corresponding to Leiden, Infomap, SBM, and Seurat, respectively. The zoomed-in figure in green highlights the performance stability across a Markov time range of 0.2 to 5. The same stability is not observed for other methods (see Figure S5). This further confirms that Infomap not only recovers the original cell types better than the other methods, but it also maintains stability and robustness over a wide range of Markov times.

**Fig. 4.**
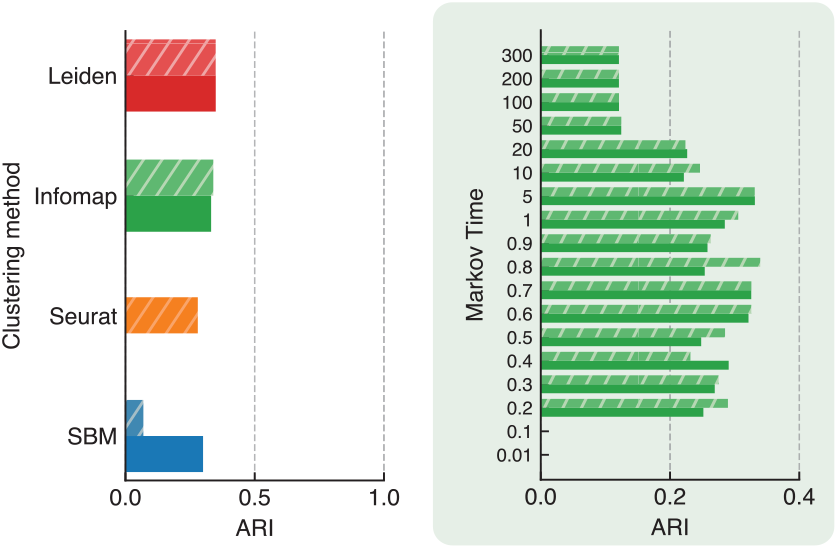
Adjusted Rand Index (ARI) between cell types and detected clusters for SBM, Seurat, Infomap, and Leiden in the 68k PBMCs full dataset. WGCNA and scGNN are not included due to their high time complexity. Both the weighted and unweighted versions of the networks were considered for algorithms that can handle both. The zoomed-in figure in green illustrates the ARI for Infomap across different Markov times.

### 3.2. Results on PBMCs: Alternative preprocessing

In this section we explore different combinations of preprocessing steps to generate networks and then apply the clustering algorithms. The aim is to identify pipelines that lead to network representations whose topology makes cell types more visible. We also compare the results acquired from the networks generated by Seurat’s pipeline with the ones obtained from the alternate pipelines we explored. In Figure 5 we show the clusters found by each clustering algorithm on network generated from the full 68k dataset by Seurat’s pipeline (top row) and the network generated using the alternative pipeline with the highest correspondence between cell types and detected partitions (bottom row). The alternate pipeline in this case consists of KNN with 20 neighbors, normalization, log transformation, and feature selection. Although the two network structures have a similar overall structure, they exhibit considerable differences. In particular, the community structures vary significantly across the methods and network types, resulting in different Adjusted Rand Indices (ARIs) among them. These ARIs are compared in Figures 6, 7, and 8 for the three PBMC datasets.

**Fig. 5.**
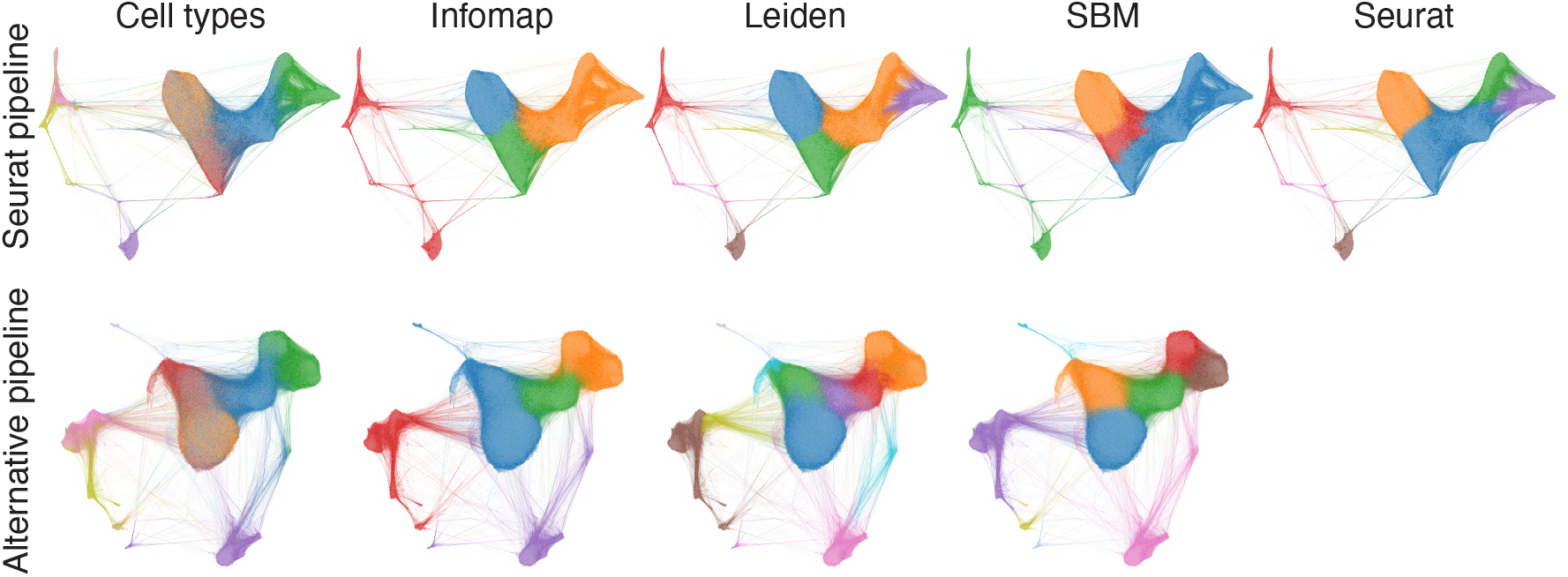
Visualizations of the generated networks from Seurat’s (top row) and alternative (bottom row) pipelines. The colors correspond to the cell types in the first column on the left, and clusters identified by the clustering techniques otherwise. The visualizations were obtained using the Force Atlas 2 algorithm [53] implemented in Helios-Web [54].

**Fig. 6.**
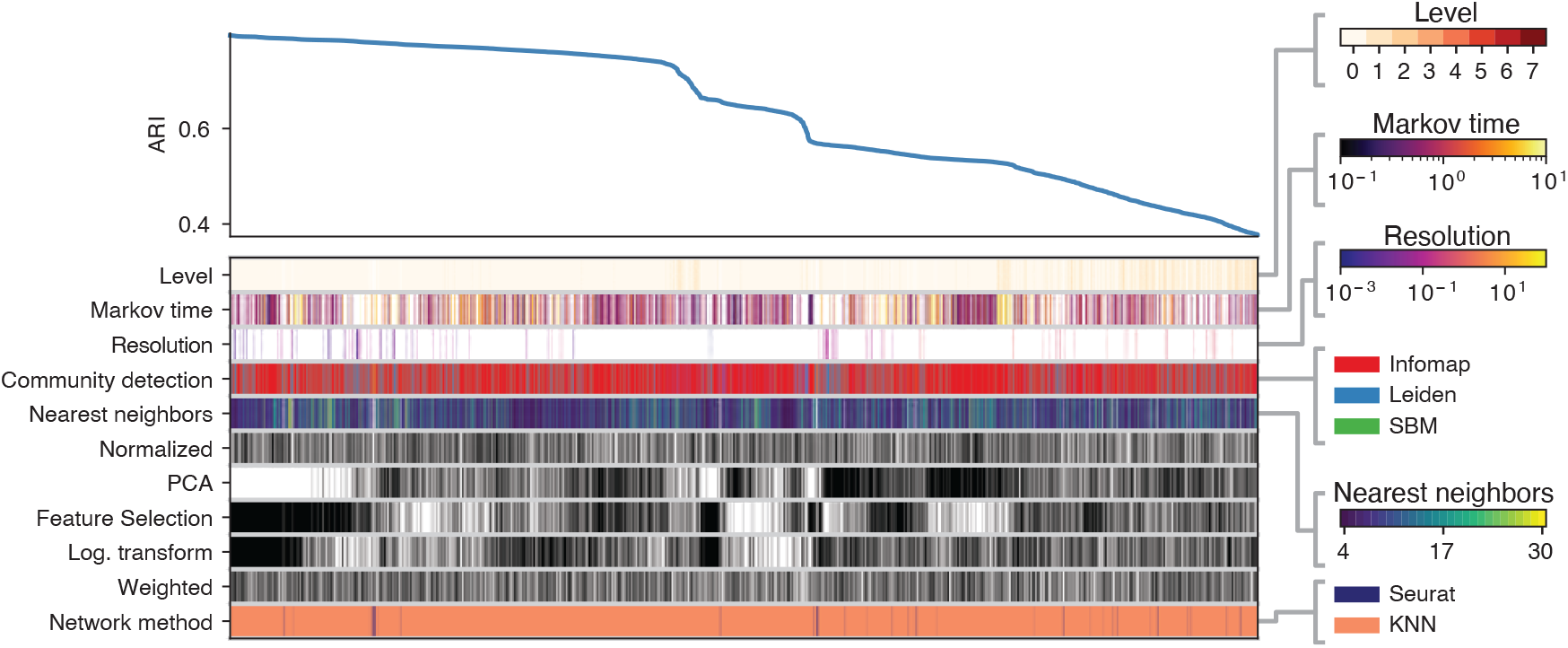
ARI across different configurations of alternative preprocessing pipelines for the 20k simple case. The top left curve reports the ARI across the top 5000 preprocessing procedures. Each column defines the specific pipeline adopted, by specifying which ingredients are present, according to the color of the blocks. For ingredients “Weighted”, “Log. transformation”, “Feature selection”, “PCA”, and “Normalization” black (white) block corresponds to having (omitting) that ingredient in the pipeline. Color bars on the right represent the value or category of the rest of the blocks in the columns.

**Fig. 7.**
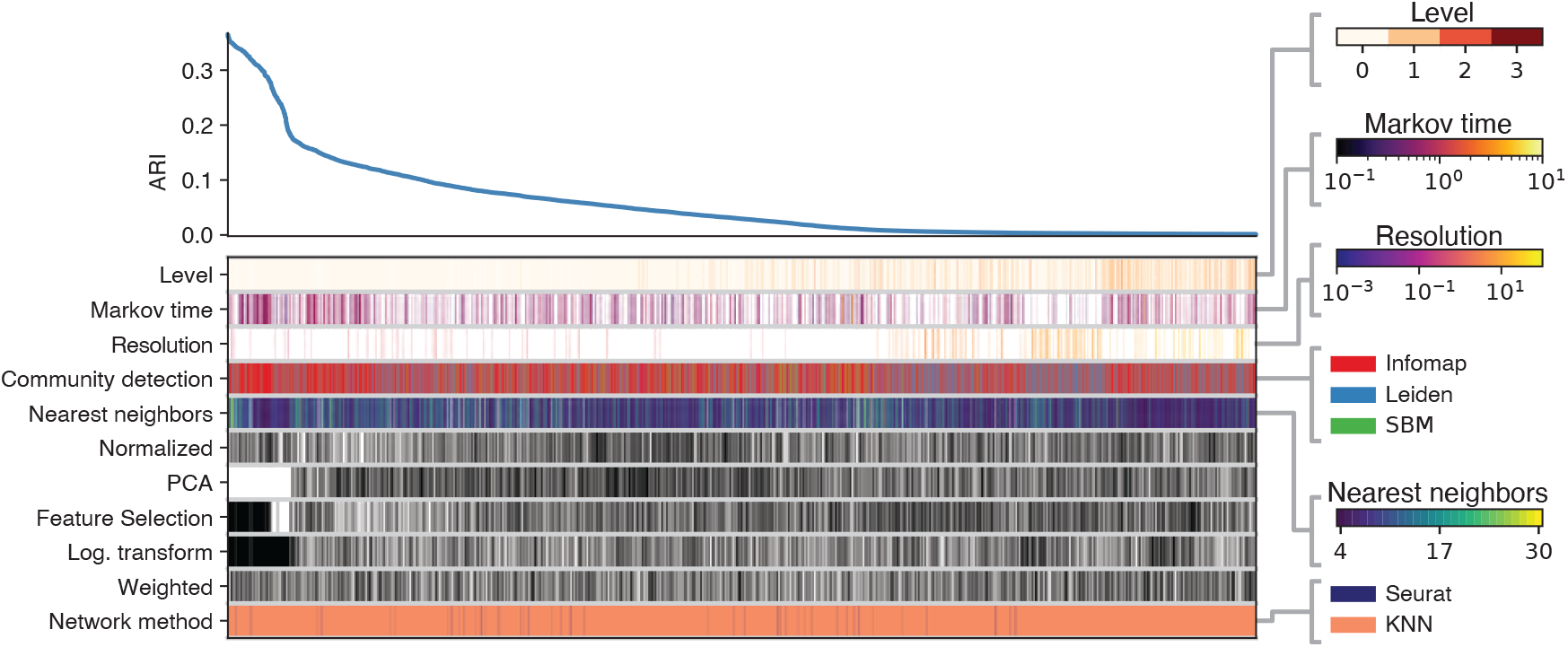
ARI across different configurations of alternative preprocessing pipelines for the 2k challenging dataset. The top left curve reports the ARI across the top 5000 preprocessing procedures. Each column defines the specific pipeline adopted, by specifying which ingredients are present, according to the color of the blocks. For ingredients “Weighted”, “Log. transformation”, “Feature selection”, “PCA”, and “Normalization” black (white) block corresponds to having (omitting) that ingredient in the pipeline. Color bars on the right represent the value or category of the rest of the blocks in the columns.

**Fig. 8.**
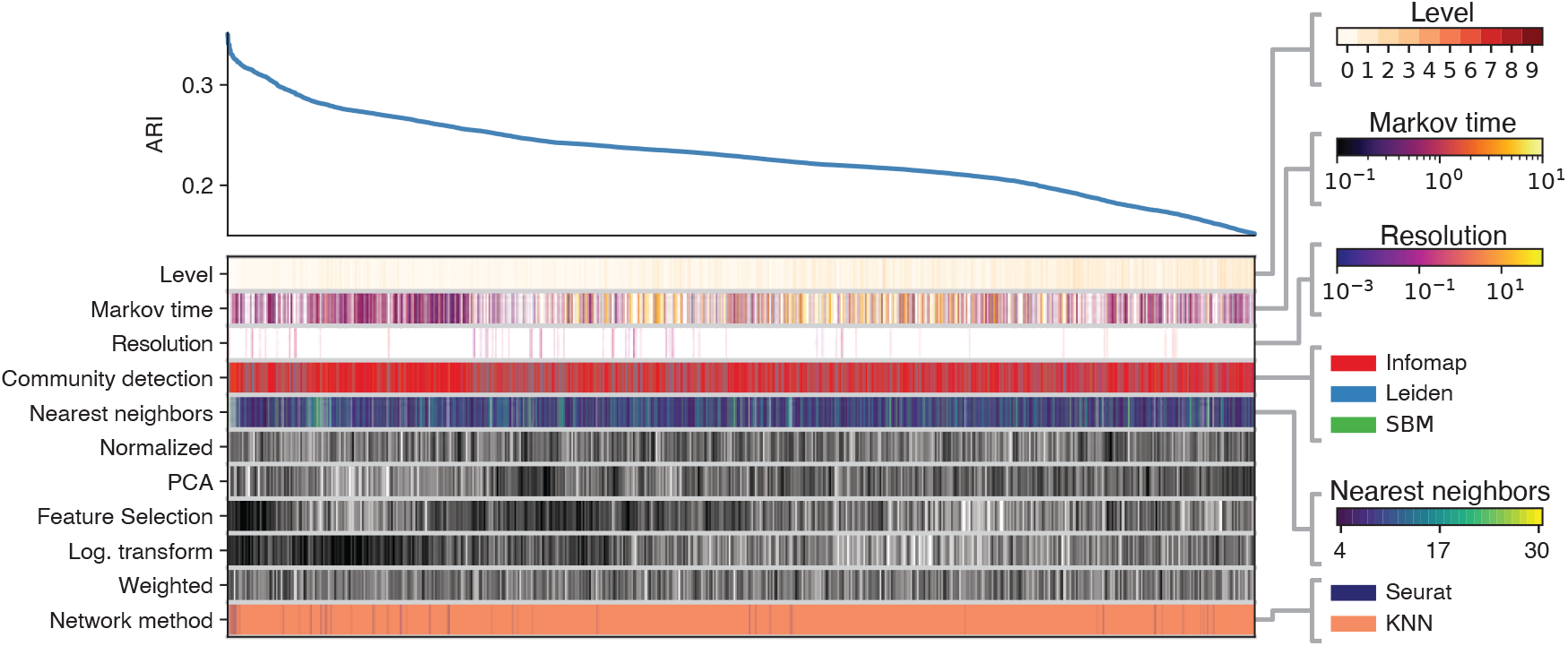
ARI across different configurations of alternative preprocessing pipelines for the 68k PBMCs full dataset. The top left curve reports the ARI across the top 5000 preprocessing procedures. Each column defines the specific pipeline adopted, by specifying which ingredients are present, according to the color of the blocks. For ingredients “Weighted”, “Log. transformation”, “Feature selection”, “PCA”, and “Normalization” black (white) block corresponds to having (omitting) that ingredient in the pipeline. Color bars on the right represent the value or category of the rest of the blocks in the columns.

Figure 6 represents the top 5000 preprocessing combinations yielding the highest ARI for the 20k dataset with highly distinct cell types. The top left curve displays the ARI across various configurations. Below it, the columns indicate the combinations of preprocessing steps and clustering algorithms used to achieve each specific ARI, with color bars on the right representing the value or category of each block in the columns. This figure demonstrates that the networks which yield the highest ARIs are generated using a KNN-based pipeline, feature selection, and log transformation, while PCA was not utilized. Additionally, the highest ARI values for the alternate pipeline exceeded 0.795 — outperforming the results achieved using Seurat’s pipeline described in Section 3.1; KNN with a relatively small number of neighbors exhibits exceptional performance. The number of possible configurations within Seurat’s analytical pipeline is notably smaller compared to the KNN-based approach. This distinction arises because the analysis in section 3.1 is limited to variations in community detection algorithms and their associated parameters. This disparity is evident in the bottom row of Figure **??**, where the dominance of the KNN-based pipeline can be attributed to the substantially greater number of combinatorial possibilities available in the KNN-based approach compared to the Seurat pipeline. Consistent with the observations in 3.1, Infomap and Leiden do better than the rest of the clustering methods. These results also indicate that certain choices in the pipeline such as using weighted versus unweighted edges and the normalization step, do not significantly affect the outcome.

Figure 7 shows the ARIs across pipelines in the same settings for the challenging sample of the PBMC 68k dataset. Similar to the previous case, the best results were obtained when feature selection and log transformation were applied, while PCA was not. The use of weights or normalization has no significant impact on performance. In this instance, however, KNN with a higher number of neighbors (*>* 15) resulted in the highest ARI. Infomap emerges as the best-performing clustering method overall. However, in certain cases, Leiden surpasses Infomap; given that their ARI values are closely comparable and Leiden is significantly more time-efficient, Leiden may also be a good choice, depending on the specific goals of the clustering analysis.

Finally, Figure 8 presents the results of the same analysis for the full PBMC 68k dataset. Notably, Seurat’s pipeline has the highest overall performance. However, within the top 5000 scores, we found similar scores for pipelines using the KNN representation and various combinations of preprocessing steps. Interestingly, neither PCA nor normalization does not play a role in the pipeline. In contrast, feature selection and log transformation appear to play a crucial role in the performance on this dataset.

### 3.3. Results on ROSMAP: Seurat-generated networks

In this section, we present the results of applying the clustering algorithms described in section 2.3.1 to networks derived from the ROSMAP dataset, as detailed in section 2.2. We employed the preprocessing pipeline from Seurat, following the same approach outlined in section 3.1. While the dotplot in Figure 1 shows close alignment between marker gene expression levels and the assigned cell types, the latter is inferred through Seurat and computational steps. A more concrete approach to evaluate the clustering results is to use the marker gene expression data as the ground truth; in this section, we use them as discriminants. Ideally, cells are considered to belong to the same type if they all express the same marker genes. Therefore we consider the marker genes as cluster labels. However, since the same cell can express multiple marker genes, we end up with a so-called fuzzy partition here, where each cell has multiple labels. Luckily in the literature there are scores that can assess the similarity between such fuzzy partition and the hard partitions delivered by the clustering algorithms. A popular metric is the Omega index [55] (See Supplemental Material for the mathematical details). The Omega index ranges between 0 and 1; a value closer to 1 indicates a higher agreement between the two partitions.

In this dataset, cell types “OPC” and “oligodendrocytes” exhibit highly similar gene expression profiles. Cells labeled with “neuron”, “excitatory neuron” and “inhibitory neuron” also share substantial similarity in gene expression. Such overlaps introduce significant fuzziness in the ground truth, and we anticipate that the Omega index will not be particularly high across methods. Nonetheless, the goal is to identify which clustering algorithm provides the highest and most robust Omega index values, reflecting the best performance. We calculated the Omega index using the Python package developed by Murray et al. [56].

The results illustrated in Figure 9 show the Omega index achieved by each clustering method. Notably, the highest scores are attained by Infomap, Leiden, and Seurat, reaching a value of 0.29. Moreover, the zoomed-in figure in green showcases the range of Markov times (0.3 to 800) over which Infomap’s performance is stable, thus reinforcing the conclusions drawn from the analysis of PBMC data. For comparisons with other methods across different resolution parameters, see Figure S6. The SBM and WGCNA methods yield Omega indices of 0.25 and 0.23, respectively, reflecting a slight decrease in performance. However, their substantial time complexity poses a significant challenge. Leiden, Seurat, and Infomap exhibit considerably lower time complexity, thereby showing their superiority in practical applications.

**Fig. 9.**
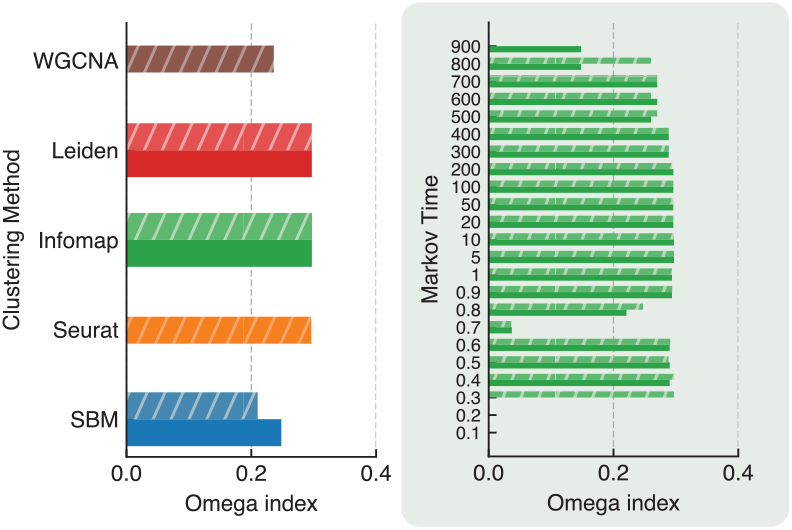
Omega index between ground truth and detected clusters for SBM, Seurat, Infomap, Leiden, and WGCNA in the ROSMAP full dataset. Both the weighted and unweighted versions of the same networks were considered for algorithms that can handle both. The zoomed-in panel illustrates the Omega index for Infomap across different Markov times.

The cell types assigned to the data also yield the same Omega index as the partitions found by three clustering algorithms, which shows that they are reliable. Indeed, in Fig. S7 we calculated the ARIs between such cell types and the partions obtained from the clustering algorithms. This figure shows the same trends observed in Figure 9. Notably, the highest ARIs are attained by Infomap, Leiden, and Seurat, reaching a value of 0.96 (see Figure S8 for the illustration of the network). This confirms the strong alignment between the assigned cell types and the clusters found by the algorithms.

Also, Infomap’s performance is stable, thus reinforcing the conclusions drawn from the prior analyses. For a comparison with other methods across different resolution parameters, see Figure S9.

### 3.4. Results on ROSMAP: Alternative preprocessing

In this last section we explore different combinations of preprocessing steps to generate networks and then apply the clustering algorithms for ROSMAP data. The setting in this investigation is the same as in section 3.2. Figure 10 represents the top 5000 preprocessing combinations in terms of ARI. Notably, the top scores are obtained using log transformation, PCA, and normalization, while feature selection appears to impact the performance negatively. Additionally, Infomap consistently stands out as the top-performing clustering method. The findings in this analysis are in line with the previous investigations in this work.

**Fig. 10.**
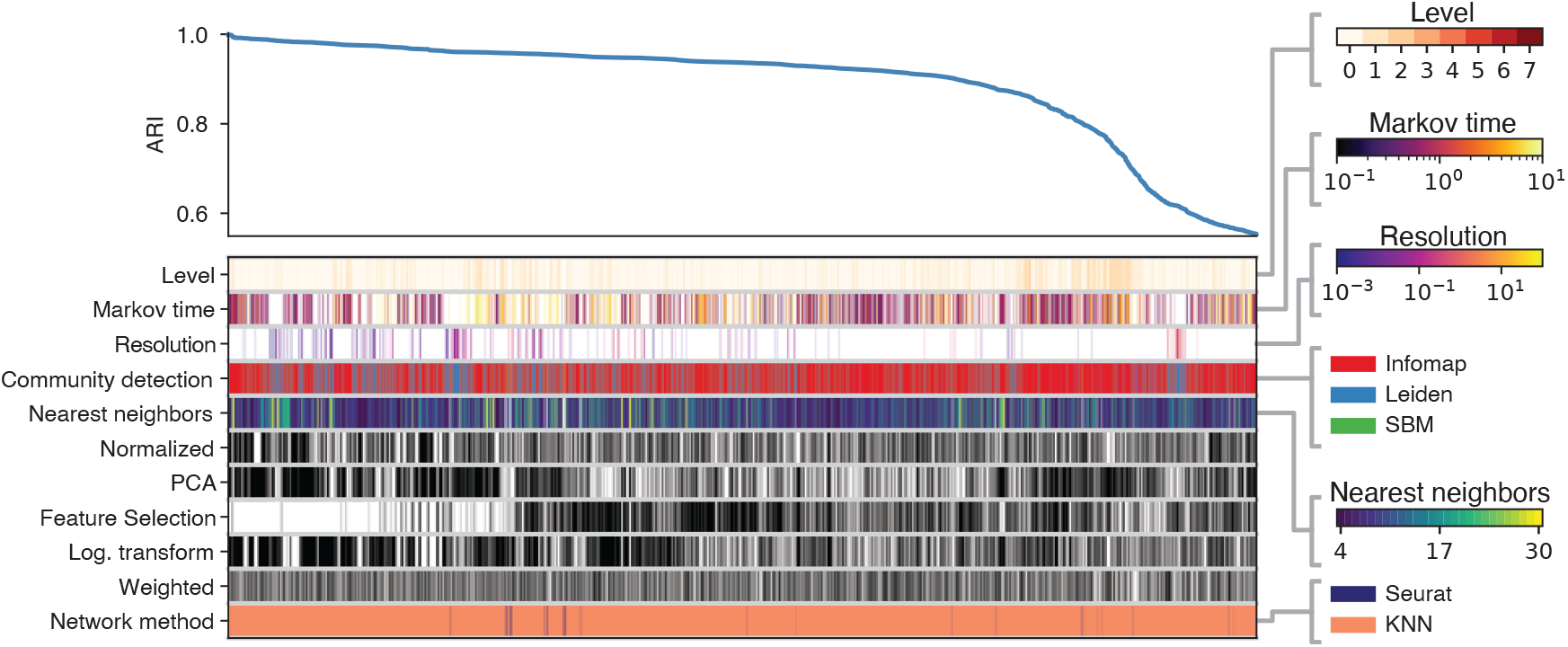
ARI across different configurations of alternative preprocessing pipelines for the ROSMAP dataset. The top left curve reports the ARI across the top 30 preprocessing procedures. Each column defines the specific pipeline adopted, by specifying which ingredients are present, according to the color of the blocks. For ingredients “Weighted”, “Log. transformation”, “Feature selection”, “PCA”, and “Normalization” black (white) block corresponds to having (omitting) that ingredient in the pipeline. Color bars on the right represent the value or category of the rest of the blocks in the columns.

## 4. Discussion and Conclusion

Single-cell RNA sequencing (scRNA-seq) analysis offers unique insights into the complexities of cellular behavior and identity. As single-cell investigations have become essential for analyzing individual cells, they enable researchers to trace developmental lineages and identify distinct cell types. This capability is particularly important in neurodegeneration and brain health research, where cell-specific dysfunction underlies the biological pathophysiology of diseases. Thus, understanding the roles of various cell types can help deepen our molecular and physiological understanding of diseases and advance therapeutic development and diagnostics.

In the context of scRNA-seq, automated procedures for identifying cell types within cell-cell networks derived from expression data are crucial. Network clustering algorithms serve as natural tools for this task; however, the literature lacks comprehensive comparative analyses of state-of-the-art methods, along with guidance on their strengths and limitations.

In this paper, we conducted a thorough comparative analysis using Seurat, Leiden, and WGCNA, alongside methods which are less familiar to biologists, like Infomap and SBM. Each method presents unique advantages and shortcomings. For instance, while Infomap demonstrates strong stability and performance, it may encounter challenges related to time complexity when applied to larger datasets. Seurat with Louvain modularity maximization is widely used and integrates seamlessly with established bioinformatics workflows, but it is not as stable as Infomap. WGCNA, although useful for constructing co-expression networks, tends to underperform in terms of ARI in this analysis, potentially due to enforcing an power law degree distribution on the network construction process. SBM provides a probabilistic framework that can be advantageous but suffers from limited resolution and higher computational costs.

Our findings indicate that Infomap consistently performs well across distinct datasets, demonstrating stability across various Markov times, and effectively identifies cell types even when they are structurally blended. The capability to adjust Markov time enables researchers to control the granularity of clustering, a crucial factor in neurodegeneration research. For instance, at shorter Markov times, Infomap captures more localized interactions, potentially uncovering distinct subpopulations of closely related cells. In Alzheimer’s disease, this can help distinguish between activation states of microglia responding to neuroinflammation or reactive astrocyte clusters with different functional roles. As Markov time increases, Infomap integrates longer range interactions within the network; this characteristic of Infomap addresses a critical gap, enabling researchers to explore a variety of cell type structures and transitions that contribute to Alzheimer’s progression.

Moreover, we highlight the critical role of preprocessing—the pipeline that transforms raw gene expression data into networks. Our dynamic platform generates multiple preprocessing pipelines combining various operations, such as log-normalization, PCA, and KNN, along with tailored parameters for specific clustering algorithms. Through extensive testing of thousands of combinations, we observed that KNN is essential for optimal performance, while steps like log-normalization, PCA, and feature selection are less critical or even irrelevant despite their common inclusion in established pipelines like Seurat. We believe this investigation will become increasingly valuable for researchers utilizing scRNA-seq data.

A key limitation in using annotations as ground truth for cluster validation, as practiced in this study, lies in the assumption that algorithmically detected structural groups align perfectly with predefined categories. This assumption does not generally hold, as studies have shown that structural clusters identified by computational methods do not necessarily correspond to annotated categories [57, 58]. However, given the demonstrated association between structural clusters and cell types in prior research, it remains reasonable to evaluate which method performs best within the specific context of cell population separation.

## 5. Key Points

- We conducted a comprehensive comparative analysis of clustering algorithms for cell type identification in single-cell RNA sequencing (scRNA-seq) data. The study utilized two datasets: the Peripheral Blood Mononuclear Cell (PBMC) dataset and the Religious Orders Study (ROS)/Rush Memory and Aging Project (MAP) dataset. The algorithms evaluated include established methods widely used in biology, such as Seurat and WGCNA, as well as algorithms less frequently applied in biological contexts, such as Infomap and Stochastic Block Models (SBM).
- Among the evaluated methods, Infomap emerged as the most reliable clustering algorithm. The Leiden algorithm, which is a variant employed by Seurat, demonstrated performance comparable to Infomap. However, WGCNA proved to be less effective in this context.
- Our findings underscore the critical importance of preprocessing steps in scRNA-seq analysis, particularly in generating cell-cell networks from raw gene expression data. Key steps such as KNN-based network construction and log-normalization were found to significantly influence clustering performance, while commonly used steps like PCA and feature selection were often less impactful. We conclude that a carefully optimized preprocessing pipeline enhances both the resolution and biological interpretability of clustering results.

## 7. Data Availability

PBMC data can be accessed from github.com/10XGenomics/single-cell-3prime-paper [5]. ROSMAP resources can be requested at radc.rush.edu and www.synpase.org.

## 8. Author Contributions

F. S. F. N. conducted the majority of the analysis. F. N. S. developed the alternative preprocessing pipelines and executed the clustering algorithms on the resulting networks. S. F. conceptualized and designed the overall analysis framework. All authors contributed to the project by providing feedback at various stages and collaboratively writing the manuscript.

## References

1. Fuchou Tang, Catalin Barbacioru, Yangzhou Wang, Ellen Nordman, Clarence Lee, Nanlan Xu, Xiaohui Wang, John Bodeau, Brian B Tuch, Asim Siddiqui, et al. mrna-seq whole-transcriptome analysis of a single cell. Nature methods, 6(5):377–382, 2009.

2. Jillian J Goetz and Jeffrey M Trimarchi. Transcriptome sequencing of single cells with smart-seq. Nature biotechnology, 30(8):763–765, 2012.

3. Xueyang Shen, Mingming Li, Kangmei Shao, Yongnan Li, and Zhaoming Ge. Post-ischemic inflammatory response in the brain: Targeting immune cell in ischemic stroke therapy. Frontiers in Molecular Neuroscience, 16:1076016, 2023.

4. Evan Z Macosko, Anindita Basu, Rahul Satija, James Nemesh, Karthik Shekhar, Melissa Goldman, Itay Tirosh, Allison R Bialas, Nolan Kamitaki, Emily M Martersteck, et al. Highly parallel genome-wide expression profiling of individual cells using nanoliter droplets. Cell, 161(5):1202–1214, 2015.

5. GXY Zheng, JM Terry, P Belgrader, P Ryvkin, ZW Bent, R Wilson, SB Ziraldo, TD Wheeler, GP McDermott, J Zhu, et al. Massively parallel digital transcriptional profiling of single cells. nat. commun. 8, 14049, 2017.

6. Alex K Shalek, Rahul Satija, Xian Adiconis, Rona S Gertner, Jellert T Gaublomme, Raktima Raychowdhury, Schraga Schwartz, Nir Yosef, Christine Malboeuf, Diana Lu, et al. Single-cell transcriptomics reveals bimodality in expression and splicing in immune cells. Nature, 498(7453):236–240, 2013.

7. Sophie Petropoulos, Daniel Edsgärd, Björn Reinius, Qiaolin Deng, Sarita Pauliina Panula, Simone Codeluppi, Alvaro Plaza Reyes, Sten Linnarsson, Rickard Sandberg, and Fredrik Lanner. Single-cell rna-seq reveals lineage and x chromosome dynamics in human preimplantation embryos. Cell, 165(4):1012–1026, 2016.

8. Cole Trapnell, Davide Cacchiarelli, Jonna Grimsby, Prapti Pokharel, Shuqiang Li, Michael Morse, Niall J Lennon, Kenneth J Livak, Tarjei S Mikkelsen, and John L Rinn. Pseudo-temporal ordering of individual cells reveals dynamics and regulators of cell fate decisions. Nature biotechnology, 32(4):381, 2014.

9. Michael JT Stubbington, Tapio Lönnberg, Valentina Proserpio, Simon Clare, Anneliese O Speak, Gordon Dougan, and Sarah A Teichmann. T cell fate and clonality inference from single-cell transcriptomes. Nature methods, 13(4):329–332, 2016.

10. Hansruedi Mathys, Jose Davila-Velderrain, Zhuyu Peng, Fan Gao, Shahin Mohammadi, Jennie Z Young, Madhvi Menon, Liang He, Fatema Abdurrob, Xueqiao Jiang, et al. Single-cell transcriptomic analysis of alzheimer’s disease. Nature, 570(7761):332–337, 2019.

11. Blue B Lake, Rizi Ai, Gwendolyn E Kaeser, Neeraj S Salathia, Yun C Yung, Rui Liu, Andre Wildberg, Derek Gao, Ho-Lim Fung, Song Chen, et al. Neuronal subtypes and diversity revealed by single-nucleus rna sequencing of the human brain. Science, 352(6293):1586–1590, 2016.

12. Peter van Galen, Volker Hovestadt, Marc H Wadsworth II, Travis K Hughes, Gabriel K Griffin, Sofia Battaglia, Julia A Verga, Jason Stephansky, Timothy J Pastika, Jennifer Lombardi Story, et al. Single-cell rna-seq reveals aml hierarchies relevant to disease progression and immunity. Cell, 176(6):1265–1281, 2019.

13. Alexander B Rosenberg, Charles M Roco, Richard A Muscat, Anna Kuchina, Paul Sample, Zizhen Yao, Lucas Gray, David J Peeler, Sumit Mukherjee, Wei Chen, et al. Split-seq reveals cell types and lineages in the developing brain and spinal cord. Science (New York, NY), 360(6385):176, 2018.

14. Saskia Freytag, Luyi Tian, Ingrid Lönnstedt, Milica Ng, and Melanie Bahlo. Comparison of clustering tools in r for medium-sized 10x genomics single-cell rna-sequencing data. F1000Research, 7, 2018.

15. Yuchen Yang, Ruth Huh, Houston W Culpepper, Yuan Lin, Michael I Love, and Yun Li. Safe-clustering: single-cell aggregated (from ensemble) clustering for single-cell rna-seq data. Bioinformatics, 35(8):1269–1277, 2019.

16. Carlos Alberto Moreira-Filho, Silvia Yumi Bando, Fernanda Bernardi Bertonha, Filipi Nascimento Silva, and Luciano da Fontoura Costa. Methods for gene co-expression network visualization and analysis. In Transcriptomics in Health and Disease, pages 143–163. Springer, 2022.

17. Rahul Satija, Jeffrey A Farrell, David Gennert, Alexander F Schier, and Aviv Regev. Spatial reconstruction of single-cell gene expression data. Nature biotechnology, 33(5):495–502, 2015.

18. Yuhan Hao, Tim Stuart, Madeline H Kowalski, Saket Choudhary, Paul Hoffman, Austin Hartman, Avi Srivastava, Gesmira Molla, Shaista Madad, Carlos Fernandez-Granda, et al. Dictionary learning for integrative, multimodal and scalable single-cell analysis. Nature biotechnology, pages 1–12, 2023.

19. Ruth Huh, Yuchen Yang, Yuchao Jiang, Yin Shen, and Yun Li. Same-clustering: S ingle-cell a ggregated clustering via m ixture model e nsemble. Nucleic acids research, 48(1):86–95, 2020.

20. Vladimir Yu Kiselev, Kristina Kirschner, Michael T Schaub, Tallulah Andrews, Andrew Yiu, Tamir Chandra, Kedar N Natarajan, Wolf Reik, Mauricio Barahona, Anthony R Green, et al. Sc3: consensus clustering of single-cell rna-seq data. Nature methods, 14(5):483–486, 2017.

21. Peijie Lin, Michael Troup, and Joshua WK Ho. Cidr: Ultrafast and accurate clustering through imputation for single-cell rna-seq data. Genome biology, 18:1–11, 2017.

22. Bo Wang, Junjie Zhu, Emma Pierson, Daniele Ramazzotti, and Serafim Batzoglou. Visualization and analysis of single-cell rna-seq data by kernel-based similarity learning. Nature methods, 14(4):414–416, 2017.

23. Santo Fortunato. Community detection in graphs. Phys. Rep., 486:75–174, 2010.

24. Santo Fortunato and Darko Hric. Community detection in networks: A user guide. Physics Reports, 659:1–44, 2016. Community detection in networks: A user guide.

25. Santo Fortunato and MEJ Newman. 20 years of network community detection. Nature Physics, 18(8):848–850, 2022.

26. Vincent A Traag, Ludo Waltman, and Nees Jan Van Eck. From louvain to leiden: guaranteeing well-connected communities. Scientific reports, 9(1):5233, 2019.

27. M. Rosvall and C. T. Bergstrom. Maps of random walks on complex networks reveal community structure. Proc. Natl. Acad. Sci. USA, 105:1118–1123, January 2008.

28. Martin Rosvall, Daniel Axelsson, and Carl T Bergstrom. The map equation. The European Physical Journal Special Topics, 178(1):13–23, 2009.

29. Peter Langfelder and Steve Horvath. Wgcna: an r package for weighted correlation network analysis. BMC bioinformatics, 9:1–13, 2008.

30. Tiago P Peixoto. Hierarchical block structures and high-resolution model selection in large networks. Physical Review X, 4(1):011047, 2014.

31. Juexin Wang, Anjun Ma, Yuzhou Chang, Jianting Gong, Yuexu Jiang, Ren Qi, Cankun Wang, Hongjun Fu, Qin Ma, and Dong Xu. scgnn is a novel graph neural network framework for single-cell rna-seq analyses. Nature communications, 12(1):1882, 2021.

32. Vincent D Blondel, Jean-Loup Guillaume, Renaud Lambiotte, and Etienne Lefebvre. Fast unfolding of communities in large networks. Journal of statistical mechanics: theory and experiment, 2008(10):P10008, 2008.

33. Masoumeh Kheirkhahzadeh, Andrea Lancichinetti, and Martin Rosvall. Efficient community detection of network flows for varying markov times and bipartite networks. Physical Review E, 93(3):032309, 2016.

34. Narges Rezaie, Farilie Reese, and Ali Mortazavi. Pywgcna: A python package for weighted gene co-expression network analysis. Bioinformatics, 39(7):btad415, 2023.

35. P. Holland, K. B. Laskey, and S. Leinhardt. Stochastic blockmodels: First steps. Soc. Netw., 5:109–137, 1983.

36. Martin Rosvall and Carl T Bergstrom. An information-theoretic framework for resolving community structure in complex networks. Proceedings of the national academy of sciences, 104(18):7327–7331, 2007.

37. Ludo Waltman and Nees Jan Van Eck. A smart local moving algorithm for large-scale modularity-based community detection. The European physical journal B, 86:1–14, 2013.

38. Charles Gawad, Winston Koh, and Stephen R Quake. Single-cell genome sequencing: current state of the science. Nature Reviews Genetics, 17(3):175–188, 2016.

39. Zhe Sun, Ting Wang, Ke Deng, Xiao-Feng Wang, Robert Lafyatis, Ying Ding, Ming Hu, and Wei Chen. Dimm-sc: a dirichlet mixture model for clustering droplet-based single cell transcriptomic data. Bioinformatics, 34(1):139–146, 2018.

40. Anael Cain, Mariko Taga, Cristin McCabe, Gilad S Green, Idan Hekselman, Charles C White, Dylan I Lee, Pallavi Gaur, Orit Rozenblatt-Rosen, Feng Zhang, et al. Multicellular communities are perturbed in the aging human brain and alzheimer’s disease. Nature neuroscience, 26(7):1267–1280, 2023.

41. David A Bennett, Aron S Buchman, Patricia A Boyle, Lisa L Barnes, Robert S Wilson, and Julie A Schneider. Religious orders study and rush memory and aging project. Journal of Alzheimer’s disease, 64(1):S161–S189, 2018.

42. Tim Stuart, Andrew Butler, Paul Hoffman, Christoph Hafemeister, Efthymia Papalexi, William M Mauck, Yuhan Hao, Marlon Stoeckius, Peter Smibert, and Rahul Satija. Comprehensive integration of single-cell data. cell, 177(7):1888–1902, 2019.

43. Gilad Sahar Green, Masashi Fujita, Hyun-Sik Yang, Mariko Taga, Anael Cain, Cristin McCabe, Natacha Comandante-Lou, Charles C White, Anna K Schmidtner, Lu Zeng, et al. Cellular communities reveal trajectories of brain ageing and alzheimer’s disease. Nature, pages 1–12, 2024.

44. Mariano I Gabitto, Kyle J Travaglini, Victoria M Rachleff, Eitan S Kaplan, Brian Long, Jeanelle Ariza, Yi Ding, Joseph T Mahoney, Nick Dee, Jeff Goldy, et al. Integrated multimodal cell atlas of alzheimer’s disease. Nature Neuroscience, pages 1–18, 2024.

45. Olaf Sporns. Structure and function of complex brain networks. Dialogues in clinical neuroscience, 15(3):247–262, 2013.

46. Malte D Luecken and Fabian J Theis. Current best practices in single-cell rna-seq analysis: a tutorial. Molecular systems biology, 15(6):e8746, 2019.

47. Constantin Ahlmann-Eltze and Wolfgang Huber. Comparison of transformations for single-cell rna-seq data. Nature Methods, 20(5):665–672, 2023.

48. Felipe L Gewers, Gustavo R Ferreira, Henrique F De Arruda, Filipi N Silva, Cesar H Comin, Diego R Amancio, and Luciano da F Costa. Principal component analysis: A natural approach to data exploration. ACM Computing Surveys (CSUR), 54(4):1–34, 2021.

49. M Àngeles Serrano, Marián Boguná, and Alessandro Vespignani. Extracting the multiscale backbone of complex weighted networks. Proceedings of the national academy of sciences, 106(16):6483–6488, 2009.

50. Wei Dong, Charikar Moses, and Kai Li. Efficient k-nearest neighbor graph construction for generic similarity measures. In Proceedings of the 20th international conference on World wide web, pages 577–586, 2011.

51. Lawrence Hubert and Phipps Arabie. partitions. J. Classif., 2(1):193–218, 1985.

52. A.-L. Barabási and R. Albert. Emergence of scaling in random networks. Science, 286:509–512, 1999.

53. Mathieu Jacomy, Tommaso Venturini, Sebastien Heymann, and Mathieu Bastian. Forceatlas2, a continuous graph layout algorithm for handy network visualization designed for the gephi software. PloS one, 9(6):e98679, 2014.

54. Filipi Nascimento Silva. Helios-web (version 0.7.9), 2023. [Computer software].

55. Linda M Collins and Clyde W Dent. Omega: A general formulation of the rand index of cluster recovery suitable for non-disjoint solutions. Multivariate behavioral research, 23(2):231–242, 1988.

56. Gabriel Murray, Giuseppe Carenini, and Raymond Ng. Using the omega index for evaluating abstractive community detection. In Proceedings of workshop on evaluation metrics and system comparison for automatic summarization, pages 10–18, 2012.

57. Darko Hric, Richard K. Darst, and Santo Fortunato. Community detection in networks: Structural communities versus ground truth. Phys. Rev. E, 90:062805, Dec 2014.

58. Leto Peel, Daniel B Larremore, and Aaron Clauset. The ground truth about metadata and community detection in networks. Science advances, 3(5):e1602548, 2017.

