## Supplemental Material for "Cell Type Differentiation Using Network Clustering Algorithms"

Supplemental Material
Cell Type Differentiation Using Network Clustering Algorithms

### 1 Dependence of ARI on different resolution parameters and Markov times - PBMCs

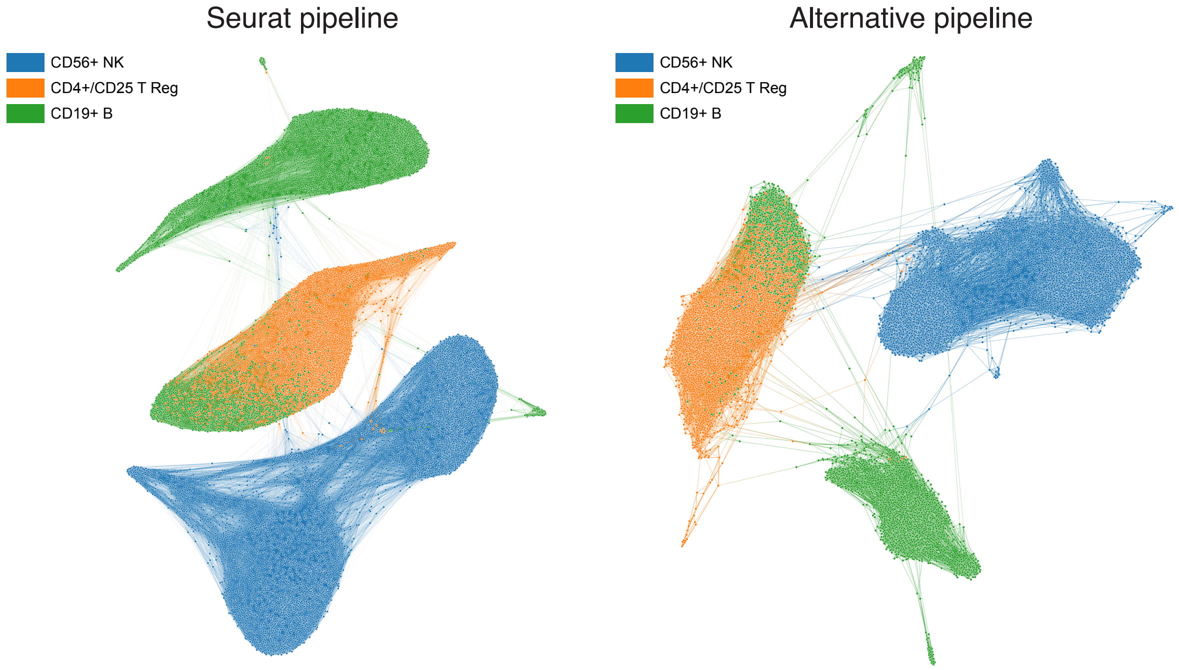


Figure S1: Illustration of the networks obtained from 20k simple PBMCs dataset. These networks are generated using Seurat and the alternative pipelines.

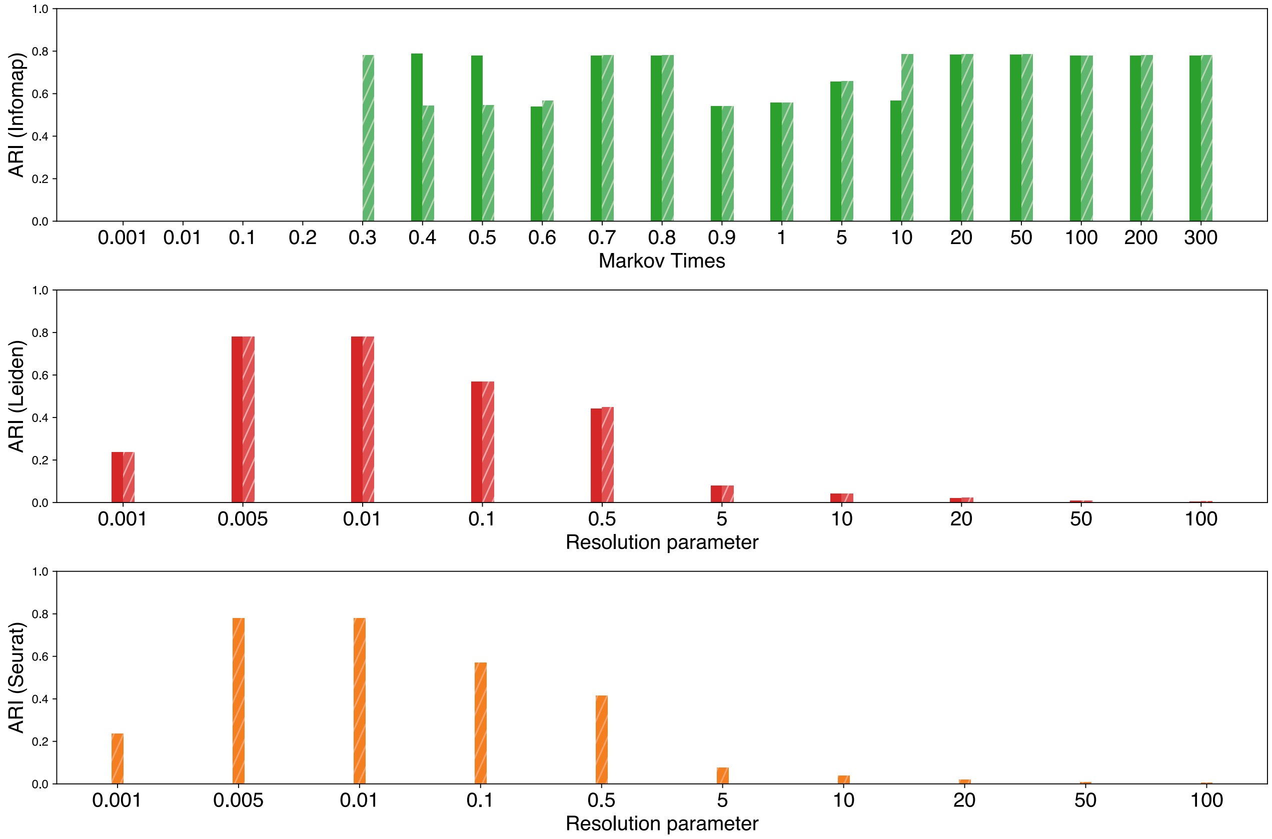


Figure S2: ARI across different resolution parameters for the network generated from 20k simple PBMCs dataset.

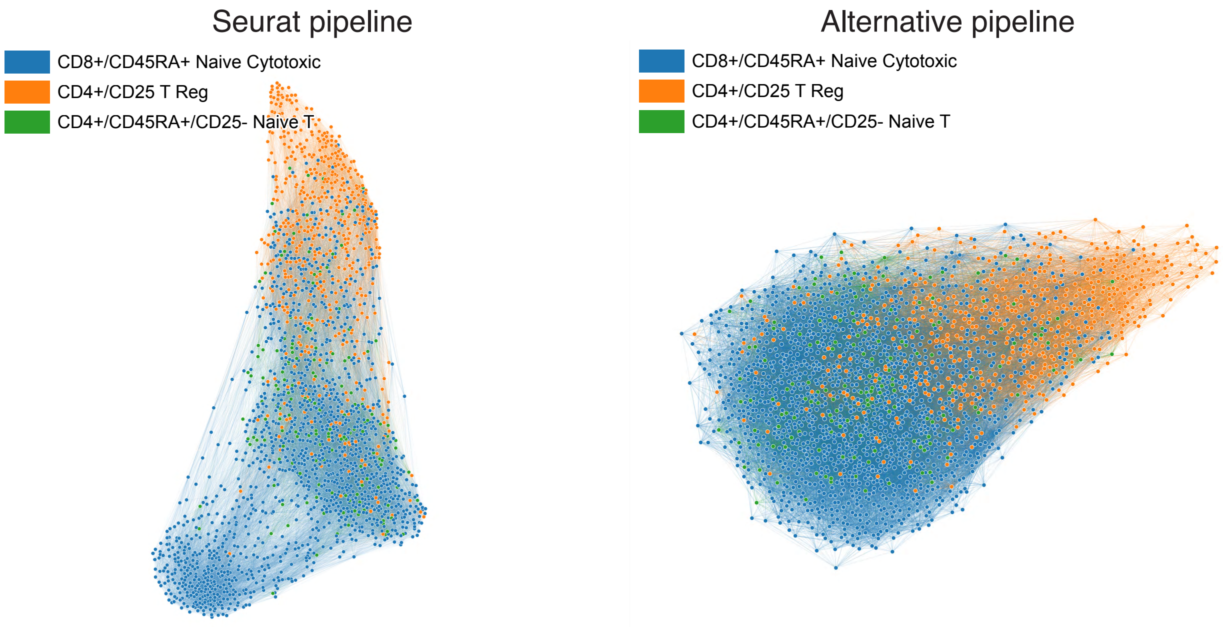


Figure S3: Illustration of the networks obtained from 2k challenging PBMCs dataset. These networks are generated using Seurat and the alternative pipelines.

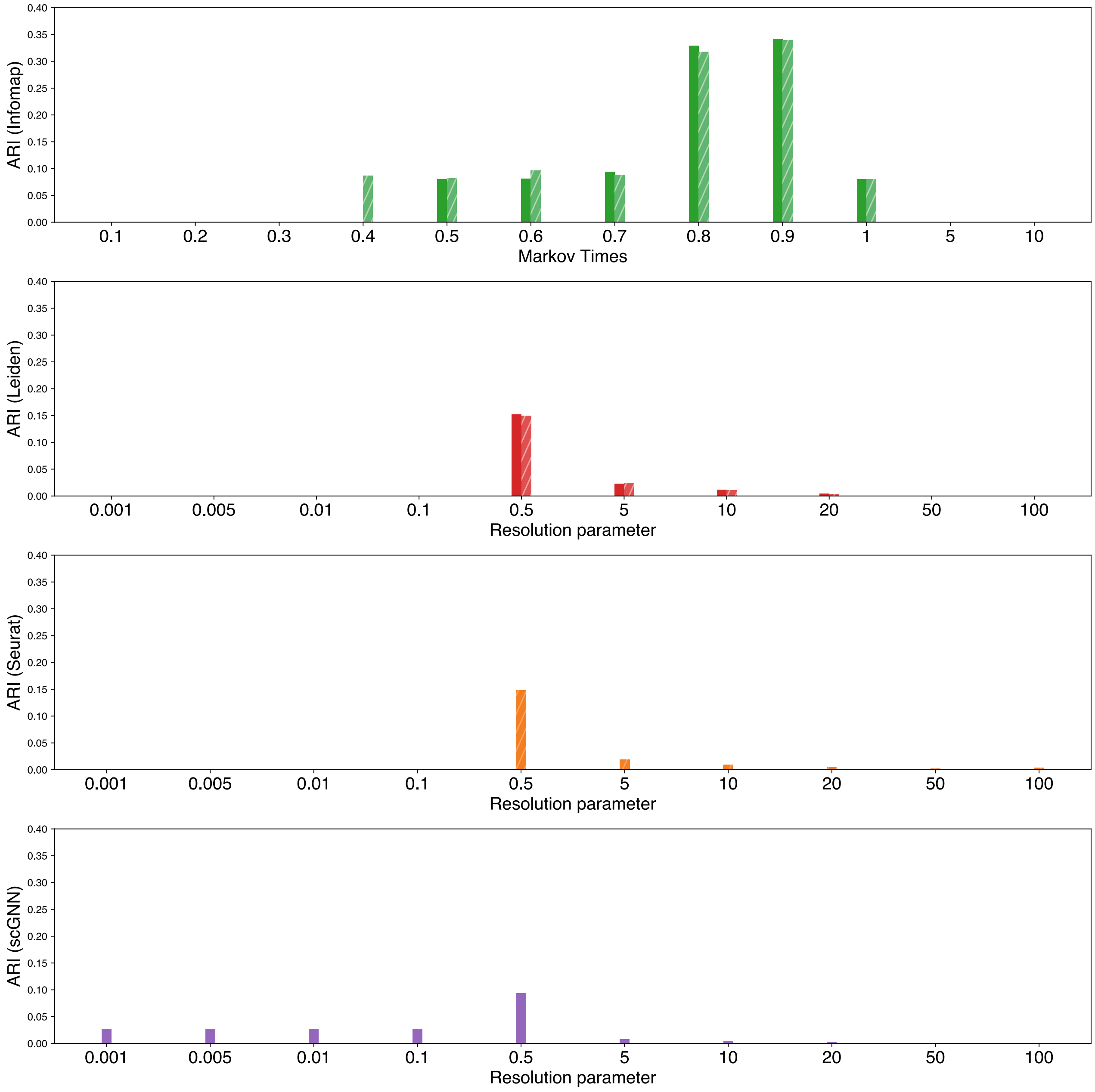


Figure S4: ARI across different resolution parameters for the network generated from 2k challenging PBMCs dataset.

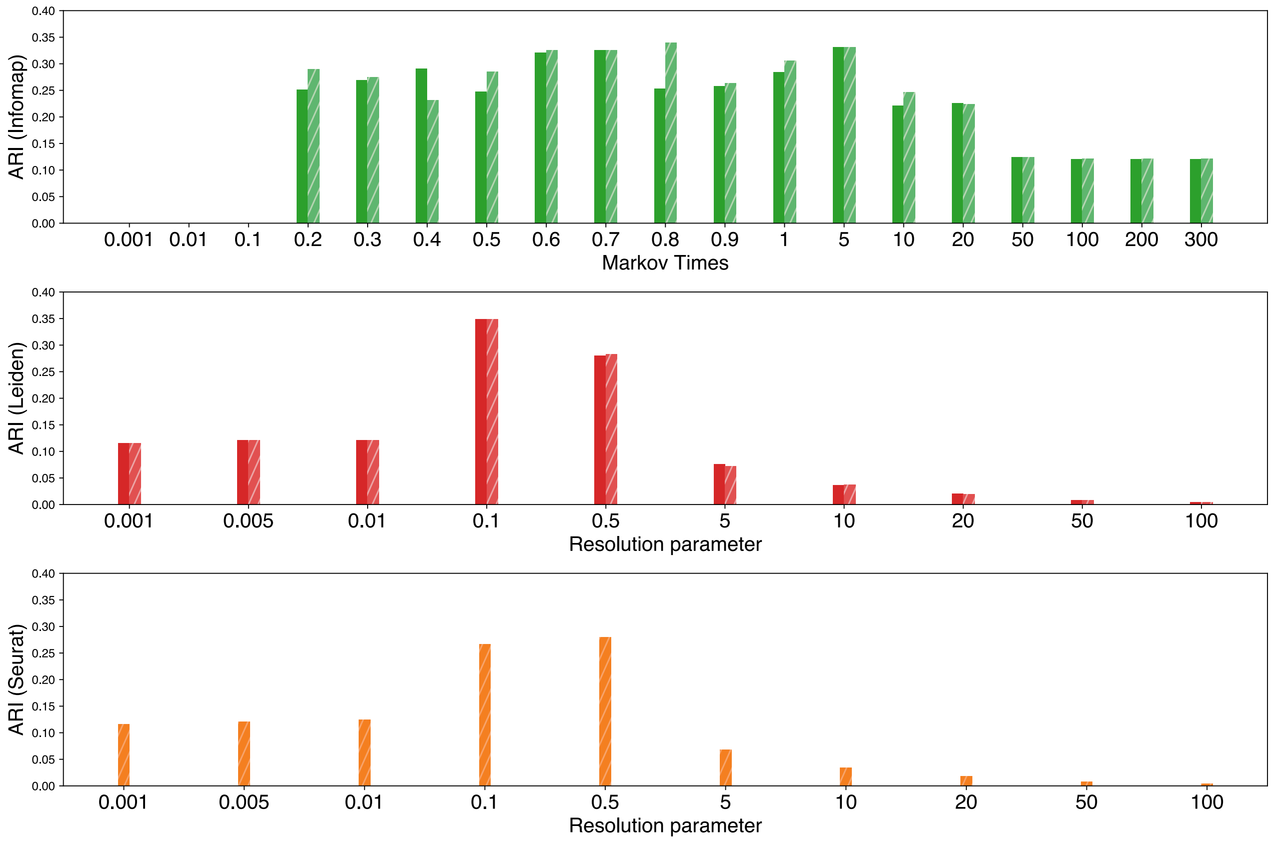


Figure S5: ARI across different resolution parameters or markov times for the Seurat-generated network from 68k PBMCs full dataset.

#### 2 Omega Index

The Omega Index $\Omega(s_1, s_2)$ measures the similarity between two clustering solutions that may include overlapping clusters. To calculate it, we first determine the observed agreement, $\text{Obs}(s_1, s_2)$, by summing the proportion of pairs that both clustering solutions agree to assign to the same number of clusters. This is expressed as

$$\text{Obs}(s_1, s_2) = \sum_{j=0}^{\min(J,K)} \frac{A_j}{N}$$

where $A_j$ is the count of pairs that both solutions agree to assign to $j$ clusters, and $N$ is the total number of pairs. We then calculate the expected agreement, $\text{Exp}(s_1, s_2)$, as

$$\text{Exp}(s_1, s_2) = \sum_{j=0}^{\min(J,K)} \frac{N_{j1} \cdot N_{j2}}{N^2}$$

where $N_{j1}$ and $N_{j2}$ represent the total pairs assigned to $j$ clusters in solutions 1 and 2, respectively. Finally, the Omega Index is calculated as

$$\Omega(s_1, s_2) = \frac{\text{Obs}(s_1, s_2) - \text{Exp}(s_1, s_2)}{1 - \text{Exp}(s_1, s_2)}$$

This index ranges from 0 to 1, with 1 indicating perfect agreement between the clustering solutions.

### 3 Dependence of ARI on different resolution parameters and Markov times - ROSMAP

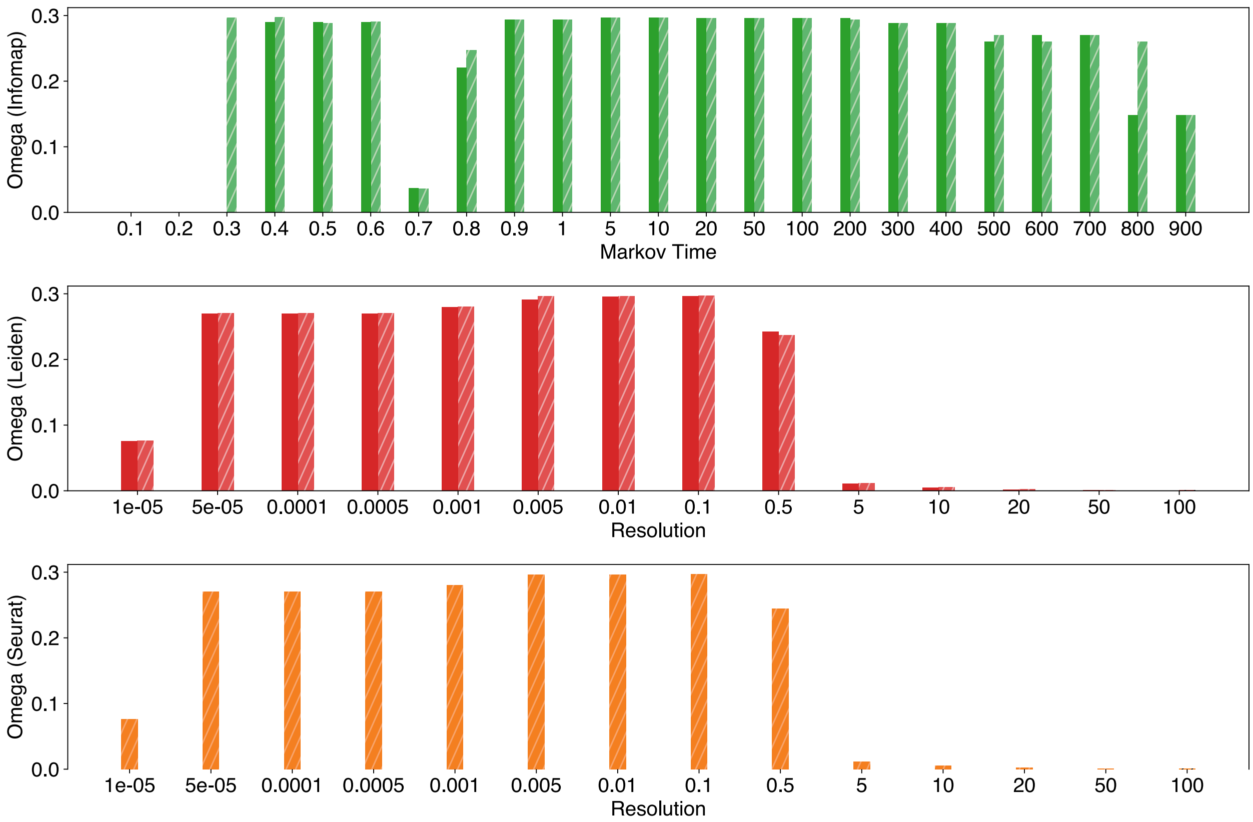


Figure S6: Omega index across different resolution parameters or markov times for the seurat-generated network from ROSMAP full dataset.

For completeness of our analysis, we calculated the ARIs illustrated in Figure S7. Notably, the highest ARIs are attained by Infomap, Leiden, and Seurat, reaching a value of 0.96. Moreover, the zoomed-in figure in green showcases the range of Markov times (0.3 to 800) over which Infomap's performance maintains stability and high efficacy, thus reinforcing the conclusions drawn from the analysis of PBMC data. For comparison with other methods across different resolution parameters, see Figure S9. The SBM and WGCNA methods yield ARIs of 0.89 and 0.85, respectively, reflecting a slight decrease in performance, yet still within a strong range. However, their substantial time complexity poses a significant challenge. Leiden, Seurat, and Infomap exhibit Considerably lower time complexity, thereby showing their superiority in practical applications.

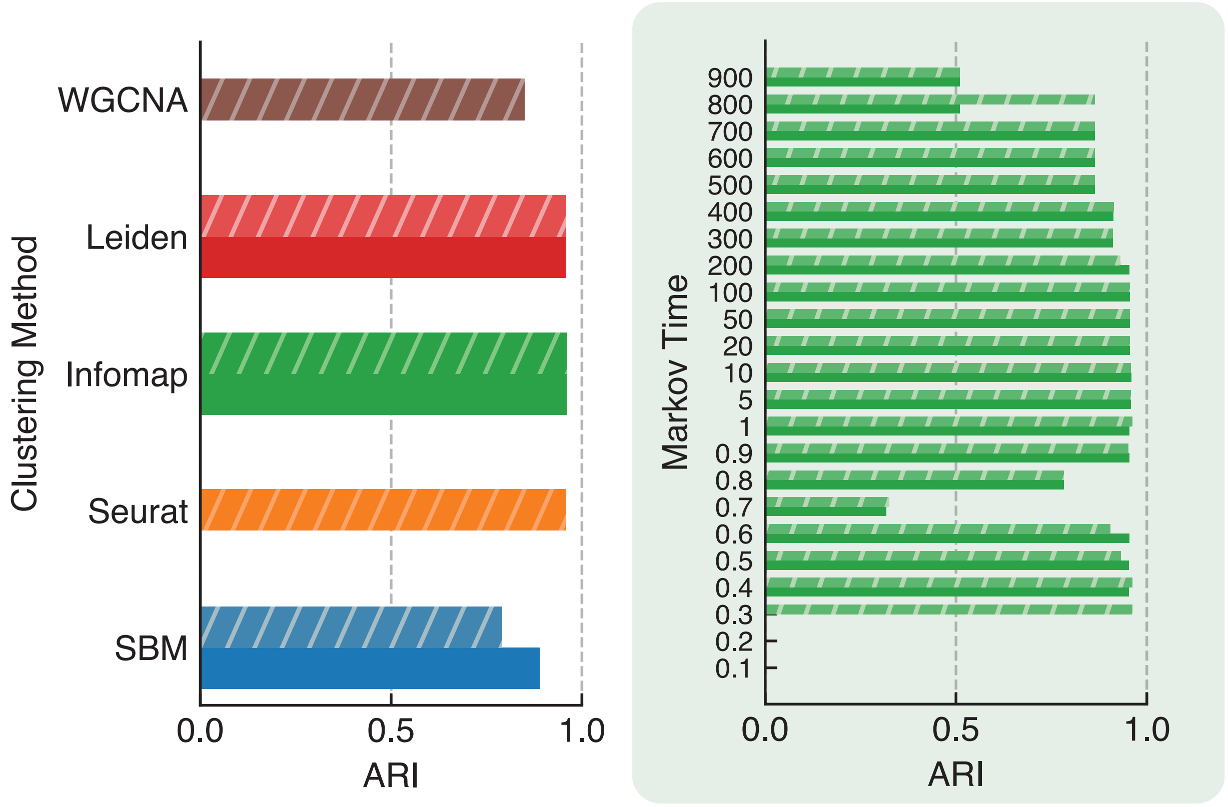


Figure S7: Adjusted Rand Index (ARI) between cell types and detected clusters for SBM, Seurat, Infomap, Leiden, and WGCNA in the ROSMAP full dataset. Both the weighted and unweighted versions of the same networks were considered for algorithms that can handle both.The zoomed-in panel illustrates the ARI across different Markov times using Infomap.

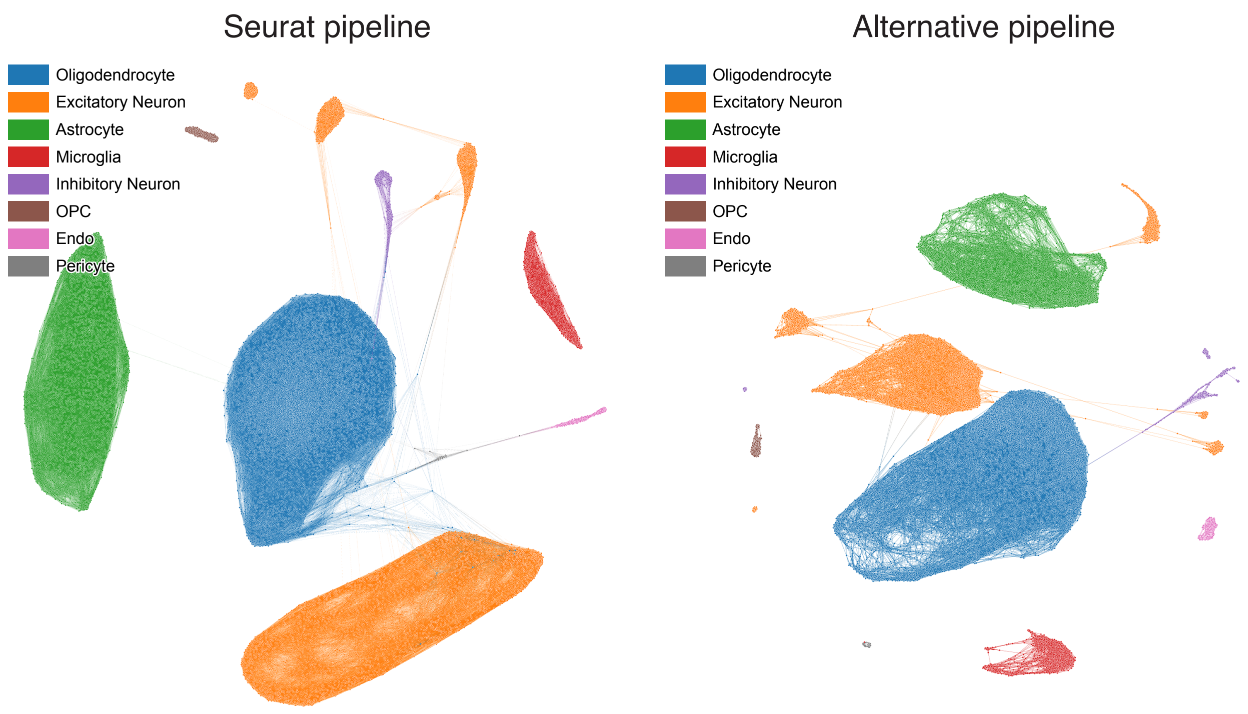


Figure S8: Illustration of the networks obtained from ROSMAP full dataset. These networks are generated using Seurat and the alternative pipelines.

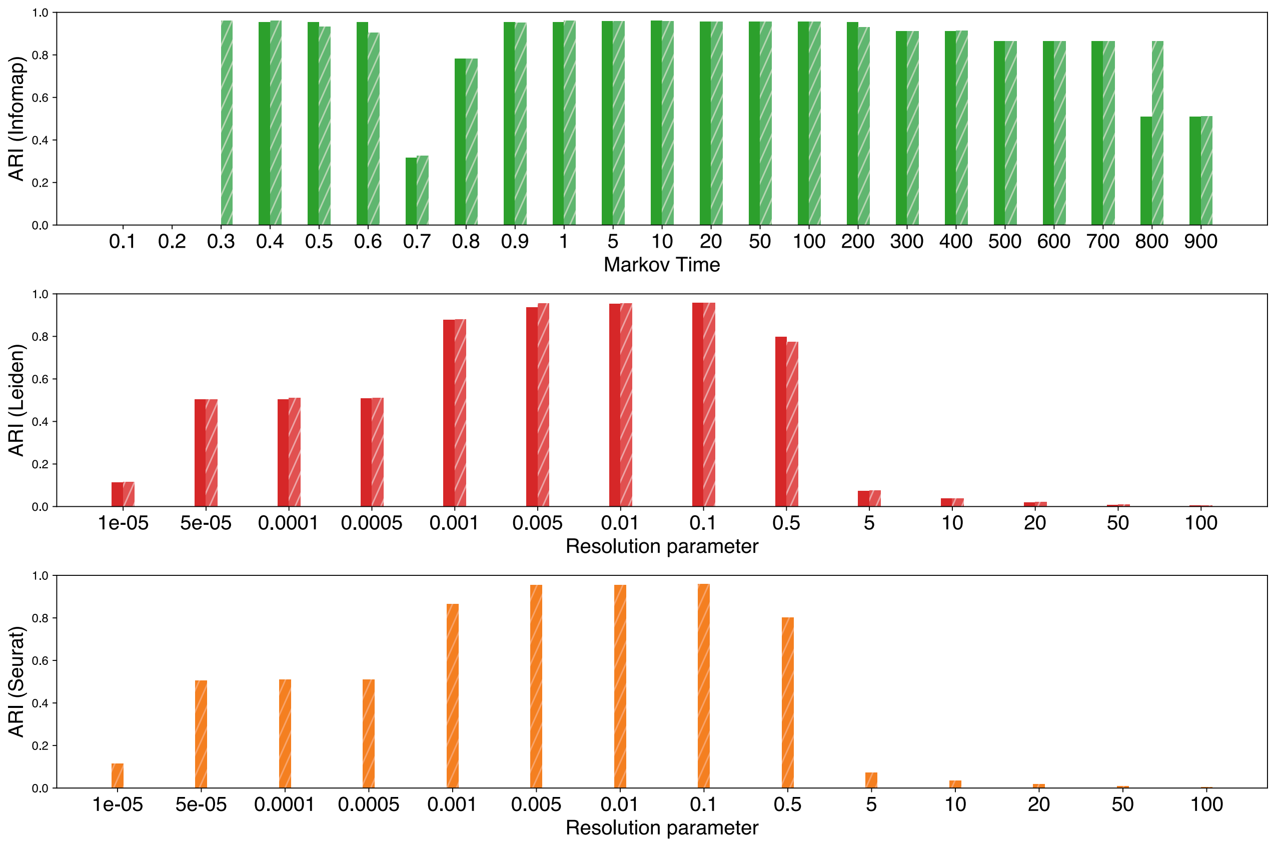


Figure S9: ARI across different resolution parameters for the network generated from ROSMAP dataset.
